# Identification of Adomavirus Virion Proteins

**DOI:** 10.1101/341131

**Authors:** Nicole L. Welch, Michael J. Tisza, Gabriel J. Starrett, Anna K. Belford, Diana V. Pastrana, Yuk-Ying S. Pang, John T. Schiller, Ping An, Paul G. Cantalupo, James M. Pipas, Samantha Koda, Kuttichantran Subramaniam, Thomas B. Waltzek, Chao Bian, Qiong Shi, Zhiqiang Ruan, Terry Fei Fan Ng, Christopher B. Buck

## Abstract

Adenoviruses, papillomaviruses, and polyomaviruses are collectively known as small DNA tumor viruses. Although it has long been recognized that small DNA tumor virus oncoproteins and capsid proteins show a variety of structural and functional similarities, it is unclear whether these similarities reflect descent from a common ancestor, convergent evolution, horizontal gene transfer among virus lineages, or acquisition of genes from host cells. Here, we report the discovery of a dozen new members of an emerging virus family, the *Adomaviridae*, that unite a papillomavirus/polyomavirus-like replicase gene with an adenovirus-like virion maturational protease. Adomaviruses were initially discovered in a lethal disease outbreak among endangered Japanese eels. New adomavirus genomes were found in additional commercially important fish species, such as tilapia, as well as in reptiles. The search for adomavirus sequences also revealed an additional candidate virus family, which we refer to as xenomaviruses, in mollusk datasets. Analysis of native adomavirus virions and expression of recombinant proteins showed that the virion structural proteins of adomaviruses are homologous to those of both adenoviruses and another emerging animal virus family called adintoviruses. The results pave the way toward development of vaccines against adomaviruses and suggest a framework that ties small DNA tumor viruses into a shared evolutionary history.

**Author Summary:** In contrast to cellular organisms, viruses do not encode any universally conserved genes. Even within a given family of viruses, the amino acid sequences encoded by homologous genes can diverge to the point of unrecognizability. Although members of an emerging virus family, the *Adomaviridae*, encode replicative DNA helicase proteins that are recognizably similar to those of polyomaviruses and papillomaviruses, the functions of other adomavirus genes have been difficult to identify. Using a combination of laboratory and bioinformatic approaches, we identify the adomavirus virion structural proteins. The results link adomavirus virion protein operons to those of other midsize non-enveloped DNA viruses, including adenoviruses and adintoviruses.

## Introduction

Polyomaviruses, papillomaviruses, and adenoviruses are historically defined as small DNA tumor viruses (Pipas 2019). Although members of a fourth animal-tropic non-enveloped DNA virus family, the *Parvoviridae*, are not known to cause tumors they share a number of biological features with traditional small DNA tumor viruses. Each of the four virus families encodes non-enveloped virion proteins with similar pentameric single-β-jellyroll core folds and members of each of the four families express functionally similar oncogenes that inactivate cellular tumor suppressor proteins (Figure 1)(de Souza, Iyer et al. 2010, Krupovic and Koonin 2017). An emerging group of animal viruses called adintoviruses appears to represent a candidate fifth family with similarities to small DNA tumor viruses https://www.biorxiv.org/content/10.1101/697771v3.

**Figure 1:**
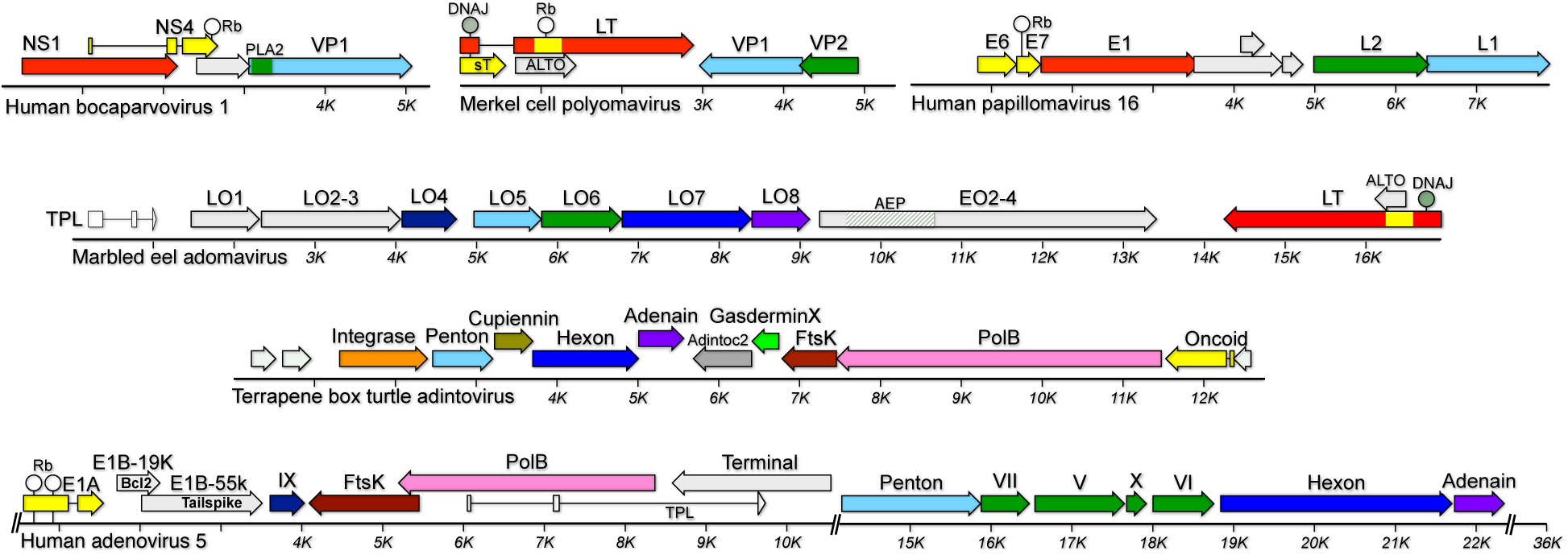
Maps of representative virus genomes. Genes are color-coded based on known or inferred functions. Polyomaviruses, papillomaviruses, and adomaviruses have circular double-stranded DNA genomes that were linearized for display. Parvoviruses have linear single-stranded DNA (ssDNA) genomes, while adintoviruses and adenoviruses have linear dsDNA genomes. Symbols and abbreviations: white lollipop, retinoblastoma/pocket protein (Rb) interaction motif; gray lollipop, domain with predicted fold similar to DNAJ chaperone proteins; AEP, domain resembling archael-eukaryotic primase small catalytic subunit; PLA2, domain with predicted fold similar to phospholipase A2. See main text for other gene names.

Polyomaviruses, papillomaviruses, and parvoviruses are proposed to have descended from circular Rep-encoding single-stranded DNA (CRESS) virus ancestors (Koonin, Dolja et al. 2015). This model explains the phylogenetic relationships of the replicative superfamily 3 helicase (S3H) and rolling circle “nickase” endonuclease domain of CRESS virus and small DNA tumor virus replicase genes (Kazlauskas, Varsani et al. 2019), but it does not account for possible similarities between the virion proteins and accessory genes of the “-oma” families and adenoviruses. Achieving a better understanding of the relationships between small DNA tumor virus families has the potential to guide comparative studies of these common human pathogens.

In 2011, a previously unknown circular dsDNA virus was discovered in a lethal disease outbreak among Japanese eels (*Anguilla japonica*)(Mizutani, Sayama et al. 2011, Okazaki, Yasumoto et al. 2016). Two related viruses have since been isolated from Taiwanese marbled eels (*Anguilla marmorata*) and a giant guitarfish (*Rhynchobatus djiddensis*)(Wen, Chen et al. 2015, Dill, Camus et al. 2018). In contrast to the eel viruses, which encode S3H replicase proteins that closely resemble the large tumor antigens (LT) of fish polyomaviruses, the guitarfish virus encodes a distinct S3H replicase, called EO1, that is distant from LT and is instead more closely related to the E1 replicases of papillomaviruses. The name “adomaviruses” has been applied to this emerging family, connoting the fact that the three known species each encode homologs of the adenain virion-maturational proteases of *ad*enoviruses as well as poly*oma*virus and papill*oma*virus S3H homologs.

The primary goal of this study is to identify the adomavirus virion proteins and to uncover possible evolutionary relationships to the virion proteins of small DNA tumor viruses. To discover additional adomavirus species, we conducted metagenomic sequencing studies and developed a pipeline to detect small DNA tumor virus-related sequences in the NCBI Sequence Read Archive (SRA). Bioinformatic methods were used to predict which adomavirus ORFs might represent virion proteins and the predictions were confirmed through functional expression in cell culture. The results pave the way toward development of preventive vaccines against pathogenic adomaviruses.

## Results

### Detection of additional adomaviruses

A post-mortem metagenomics analysis of an aquarium-bred Amazon red discus cichlid (*Symphysodon discus*) exhibiting lethargy and inflamed skin lesions revealed a complete circular adomavirus genome (Figure 2). Histopathological analysis of skin lesions from the discus specimen showed no evidence of intranuclear inclusions or other obvious histopathology.

**Figure 2:**
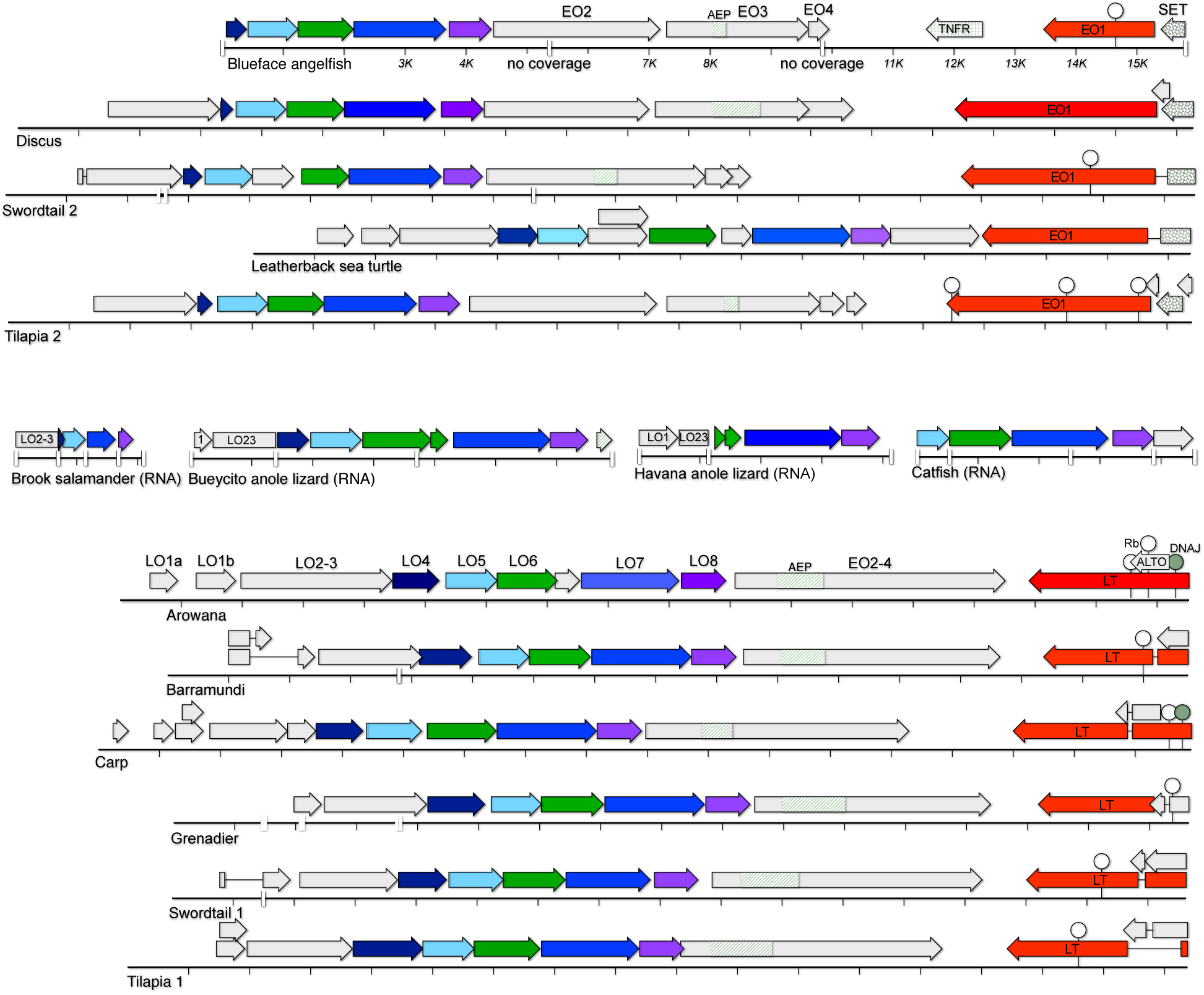
Genome maps of previously unknown adomaviruses. Genes were named based on conventions originally developed for Japanese eel adomavirus (NC_015123). Alpha adomaviruses (top half of figure) are defined by the presence of an adomavirus-specific S3H replicase, designated EO1. A row of fragmentary adomavirus sequences from RNA datasets are shown in the middle of the figure. Beta adomaviruses (bottom half of figure) encode a polyomavirus-like large tumor antigen (LT). In some cases, repetitive or GC-rich patches (particularly near the 3’ end of the LO2-3 ORF) could not be resolved using available short-read datasets. Coverage gaps are represented as white bars on the ruler. A poorly conserved set of accessory genes upstream of Alpha adomavirus EO1 genes show varying degrees of similarity to the S-adenosyl methionine-binding pocket of cellular SET proteins, which function as histone lysine-methyltransferases. Adomavirus SET homologs are highly divergent from all previously described eukaryotic and viral SET genes. The same is true for adomavirus EO2-4 and EO3 segments that encode homologs of the catalytic small subunit of archaeal eukaryotic primases (AEPs, hatched boxes). Figure supplement 1: a table of accession numbers and Linnaean designations of hosts Figure supplement 2: examples of the annotation process Figure supplement 3: GenBank-format nucleotide maps of adomaviruses Figure supplement 4: protein compilations in fasta format Figure supplement 5: splicing of marbled eel adomavirus transcripts

An adomavirus from an apparently healthy green arowana (*Scleropages formosus*) was first identified in TBLASTN searches of the NCBI Whole-Genome Shotgun (WGS) database as a set of short contigs with similarity to Japanese eel adomavirus proteins. A complete adomavirus genome was characterized by Sanger sequencing of overlapping PCR products using DNA left over from the original fin snip used for the WGS project (Bian, Hu et al. 2016).

The WGS TBLASTN survey also revealed a 4 kb contig with a sequence resembling adomavirus LT in a dataset for western softhead grenadier fish (*Malacocephalus occidentalis*) and a nearly complete adomavirus genome in a dataset for a skin biopsy of a leatherback sea turtle (*Dermochelys coriacea*). Genome sequences for the two viruses were completed using parent SRA datasets.

A pipeline using DIAMOND (a faster alternative to BLASTX (Buchfink, Xie et al. 2015)) was developed to screen SRA datasets for fish, amphibians, and reptiles. SRA datasets rich in reads encoding adomavirus-like protein sequences were subjected to *de novo* assembly. This approach resulted in the identification of seven additional adomavirus genomes (Figure 2). Notably, adomavirus sequences were found in genome sequencing datasets for farmed tilapia (*Oreochromis niloticus*), which represent a $7.5 billion per year global aquaculture industry, and in the most extensively aquacultured fish in developing countries, the mirror carp (*Cyprinus carpio*)(Bacharach, Mishra et al. 2016, Belton, Little et al. 2018). Adomavirus-like fragments were also detected in transcriptomic datasets for brook salamander (*Calotriton asper*) and two closely related species of anole lizard (genus *Anolis*).

The WGS search for adomavirus-like S3H sequences also led to the discovery of a divergent class of circular DNA elements that we tentatively designate “xenomaviruses,” connoting the exotic nature of their predicted S3H and virion proteins (Figure 2 Figure supplement 2, Figure 3, and Figure 4). Intriguingly, a conserved xenomavirus ORF shows distant predicted structural similarity to the L1 and VP1 penton proteins of papillomaviruses and polyomaviruses. The inferred replicase gene of a partial xenomavirus sequence detected in a dataset for pink abalone (*Haliotis corrugata*) encodes a domain with predicted structural similarity to the nickase domain of porcine circovirus 2 (a CRESS virus). The identification of additional viruses in this class could shed light on the evolutionary interrelationships between small DNA tumor viruses and CRESS viruses.

**Figure 3:**
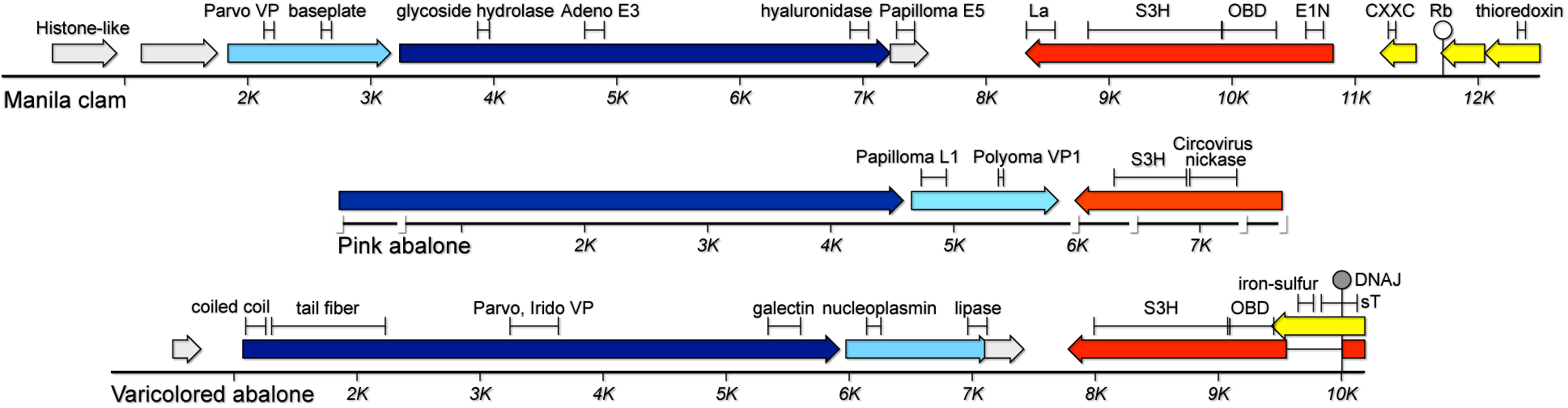
Genome maps of candidate xenomaviruses. Brackets indicate segments where remote similarities were detected in DELTA-BLAST, HHpred, and Phyre^2^ searches. The functions of these segments remain hypothetical.

**Figure 4:**
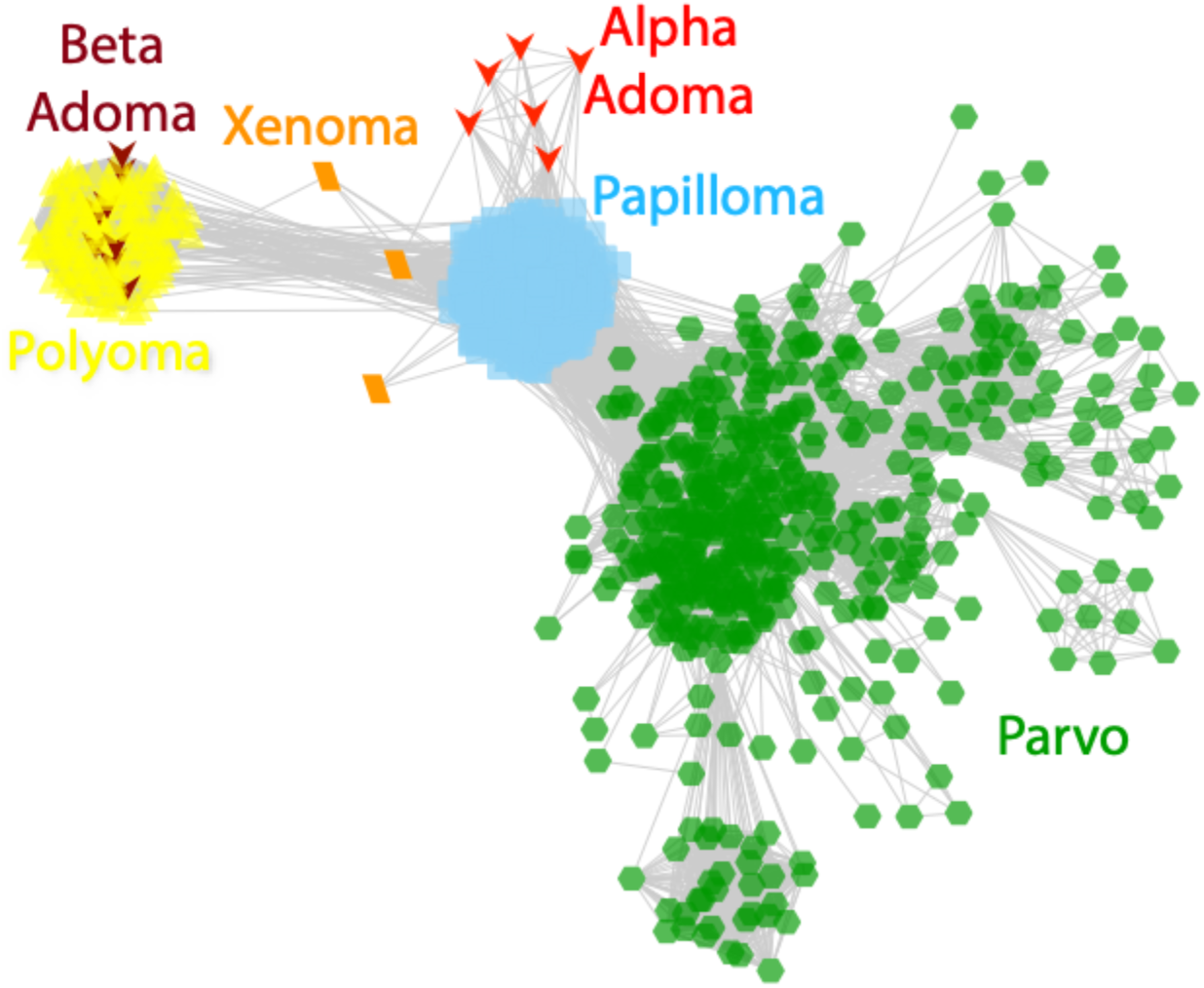
Sequence similarity network (BLASTP E-value threshold 1e-9) for S3H proteins from indicated virus groups. Figure supplement 1: an interactive version of the figure that can be viewed using Cytoscape software https://cytoscape.org Figure supplement 2: phylogenetic tree of LO8 proteins

### Adomavirus phylogeny

Adomavirus sequences can be divided into two groups based on their replicative S3H genes. A group we designate Alpha is defined by the presence of an EO1 S3H replicase gene that yields moderate hits (BLASTP E-values ∼1e-9) for papillomavirus E1 proteins. Adomavirus group Beta is defined by the presence of an S3H replicase similar (E-values ∼1e-60) to the LT proteins of polyomaviruses. A network display of BLASTP relationships is shown in Figure 4. The Alpha and Beta groupings are recapitulated in phylogenetic analyses of adomavirus LO8 (Adenain) homologs (Figure 4 Figure supplement 2).

The adomavirus LT-like proteins share a full range of familiar polyomavirus-like features, including an N-terminal DnaJ domain, a potential retinoblastoma-interaction motif (LXCXE or LXXLFD)(An, Saenz Robles et al. 2012, Gouw, Michael et al. 2018). Several Alpha adomavirus EO1 proteins encode a C-terminal domain with predicted similarity to DNA-binding RFX-type winged helices (pfam02257). The RFX-like domain is conserved at the C-terminus of S3H proteins found in larger dsDNA (for example, vaccinia virus D5 YP232992), some parvoviruses (for example, bovine parvovirus NS1 NP_041402), and xenomaviruses. HHpred analyses indicate that, like other small DNA tumor virus replicases, both classes of adomavirus S3H proteins encode a central nicking endonuclease-like origin binding domain (Hickman, Ronning et al. 2002, Iyer, Koonin et al. 2005, Koonin, Dolja et al. 2015, Kazlauskas, Varsani et al. 2019).

### Bioinformatic prediction of adomavirus virion proteins

To determine whether predicted adomavirus LO proteins are expressed from spliced or unspliced ORFs, RNAseq data published by Wen and colleagues (Wen, Chen et al. 2015) were analyzed to determine the splicing patterns of marbled eel adomavirus transcripts. Splice acceptors immediately upstream of the inferred ATG initiator codons of LO4, LO5, LO6, and LO7 proteins were extensively utilized (Figure 5 Figure supplement 5). Messenger RNAs encoding the adenovirus late genes carry a tripartite leader (TPL) that has been shown to enhance translation late in the adenovirus life cycle (Logan and Shenk 1984). A similar three-exon leader sequence was detected upstream of the marbled eel adomavirus LO genes (Figure 1).

**Figure 5:**
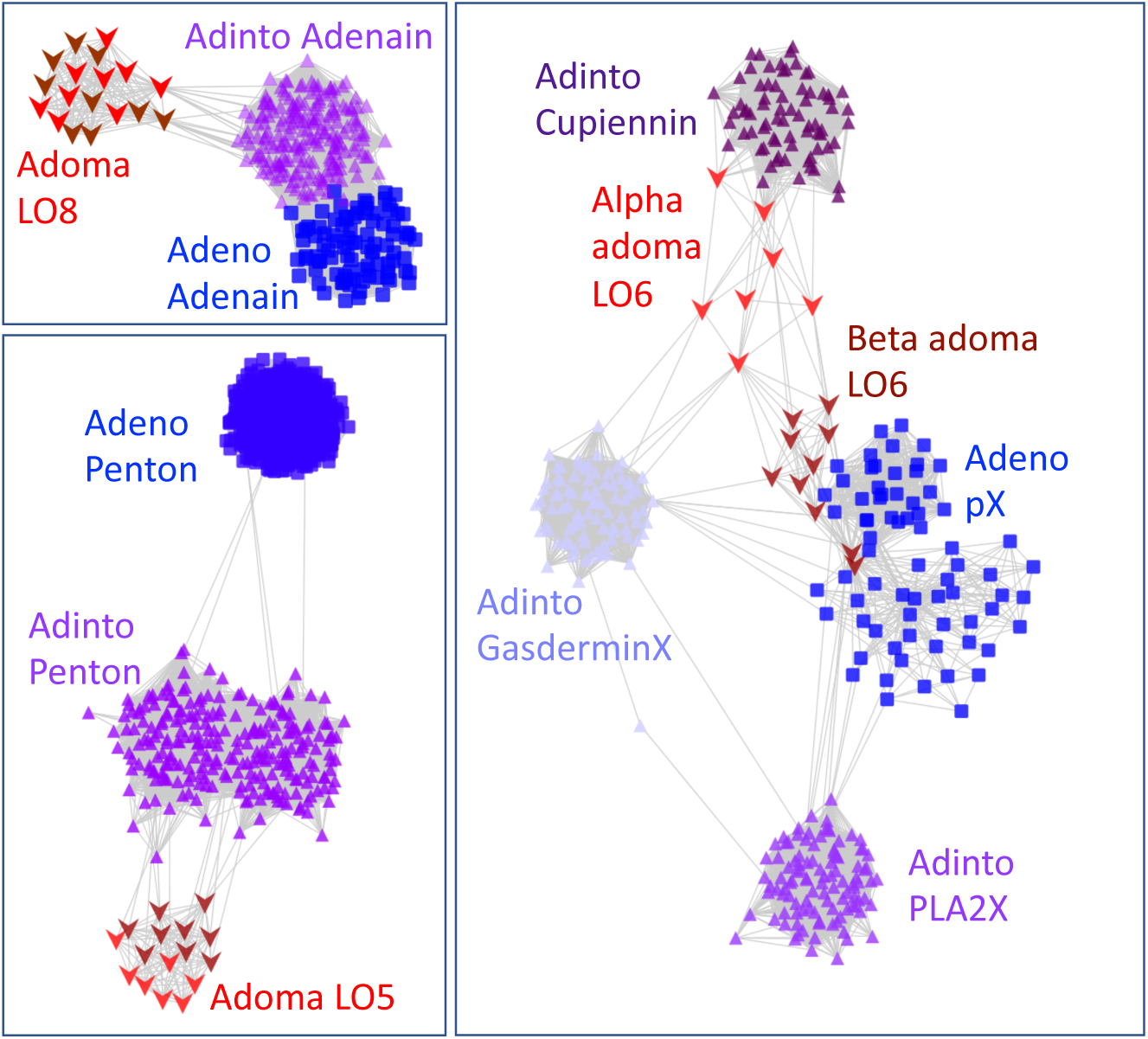
Sequence similarity network analysis of predicted minor virion proteins. The Adenain (LO8) network was constructed with a BLASTP E-value threshold of 1e-5. Penton (LO5) and virion core protein (LO6) networks used a threshold of 1e-2. Figure supplement 1: an interactive version of the figure that can be viewed using Cytoscape software https://cytoscape.org

HHpred searches were performed to detect remote similarities between the predicted structures of adomavirus proteins and known protein structures. Adomavirus LO4 proteins have a predicted C-terminal trimeric coiled-coil domain. This relatively generic predicted fold gives a large and diverse range of hits in HHpred searches, including the coiled-coil domains of various viral fiber proteins (for example, avian reovirus σC, 97%). Negative-stain EM images indicate that adomavirus virions do not have a vertex fiber (Mizutani, Sayama et al. 2011, Wen, Chen et al. 2015, Dill, Camus et al. 2018). Intriguingly, LO4 proteins yielded low probability (∼50%) HHpred hits for adenovirus pIX, a trimeric coiled coil protein that serves as a “cement” that smooths the triangular facets of the adenovirus virion. The hypothesis that LO4 is a pIX homolog could hypothetically account for the smooth appearance of adomavirus virion facets in negative-stain EM.

A C-terminal segment of some LO5 sequences, as well as alignments of multiple LO5 sequences, yielded moderate (∼60% probability) HHpred hits for CvsA1_340L protein (single-jellyroll vertex penton) of *Paramecium bursaria* chlorella virus 1 (PDB:6NCL_a6). A screen shot of a typical HHpred result is shown in Figure 2 Figure supplement 2.

Alignments of LO6 ORFs show high-probability (95%) HHPred hits for a 37 amino acid hydrophobic segment of adenovirus pX, a minor virion core protein that is thought to participate in condensation of the viral chromatin (Nemerow, Stewart et al. 2012). LO6 alignments also showed moderate probability hits (Figure 2 Figure supplement 2) for adenovirus pVI, which is believed to play a role in destabilizing cellular membranes during the infectious entry process (Moyer, Besser et al. 2015). The results suggest that LO6 might be a fused homolog of adenovirus pVI and pX virion core proteins.

HHpred searches using LO7 sequences did not produce interpretable results, with the single exception of the LO7 sequence of grenadier adomavirus, which gives a moderate-probability hit for the V20 double-jellyroll hexon major capsid protein of Sputnik virophage (Figure 2 Figure supplement 2).

In addition to offering a convenient way to summarize aggregate BLASTP interrelationships (e.g., Figure 4), all-against-all sequence similarity network analysis can be a useful method for discovering distant similarities between highly divergent groups of proteins (Iranzo, Krupovic et al. 2017). In one noteworthy example, network analyses were recently used to detect remote sequence similarities between small DNA tumor virus S3H replicases and CRESS virus replicases (Kazlauskas, Varsani et al. 2019). In contrast to traditional analyses using phylogenetic trees, it is possible for network analyses to detect individual pairs of sequences in separate clusters that both happen to have preserved the primary sequence of a common ancestor. We performed low-stringency network analyses to further investigate possible remote sequence similarities between adenovirus, adintovirus, and candidate adomavirus virion proteins.

Networks for adomavirus LO4 (inferred fiber or cement protein) and LO7 (inferred double-jellyroll hexon major capsid protein) showed few or no connections to adenovirus or adintovirus virion protein sequences, even at a BLASTP E-value threshold of 1e-1. In contrast, LO8 (adenain) proteins yielded informative networks at E-value thresholds of 1e-5 (Figure 5). At less stringent E-value thresholds (1e-2) similarities between adomavirus LO5 (inferred single-jellyroll vertex penton) and inferred adintovirus penton protein sequences emerged. PSI-BLAST searches using LO5 alignments confirmed the apparent sequence similarities (E-values ∼1e-6) to predicted adintovirus penton proteins found in arthropod and coral datasets (e.g., GBM63801 EFA12278 LSMT01002030). Although adenovirus pVI did not cluster with LO6 (inferred virion core protein) or proposed adintovirus virion core proteins (Cupiennin, GasderminX, PLA2X) at an E-value threshold of 1e-2, Alpha adomavirus LO6 proteins clustered with adintovirus cupiennin and Beta adomavirus LO6 proteins clustered with adenovirus pX proteins.

### Experimental confirmation of predicted adomavirus virion proteins

The bioinformatic results suggest that the LO4-8 operon encodes syntenic homologs of adenovirus and adintovirus virion proteins. To experimentally test this prediction, marbled eel adomavirus was grown in eel kidney cell culture (Wen, Chen et al. 2015) and virions were purified using Optiprep gradient ultracentrifugation. Virion-enriched gradient fractions were separated on SDS-PAGE gels and bands were subjected to mass spectrometric analysis (Figure 6). The analysis identified prominent bands in the Coomassie-stained gel as LO4, LO5, LO7, and LO8. The relative intensities of bands identified as LO5 and LO7 are consistent with the prediction that the two proteins constitute penton and hexon subunits, respectively. Lower molecular weight bands showed hits for LO6, suggesting that this protein is present in virions in an LO8 (adenain) cleaved form. LO6 proteins were found to encode potential adenain cleavage motifs ((MIL)XGXG or L(LR)GG) (Ruzindana-Umunyana, Imbeault et al. 2002). A list of protein modifications observed in the mass spectrometric analysis is shown in Figure 6 Figure supplement 1.

**Figure 6:**
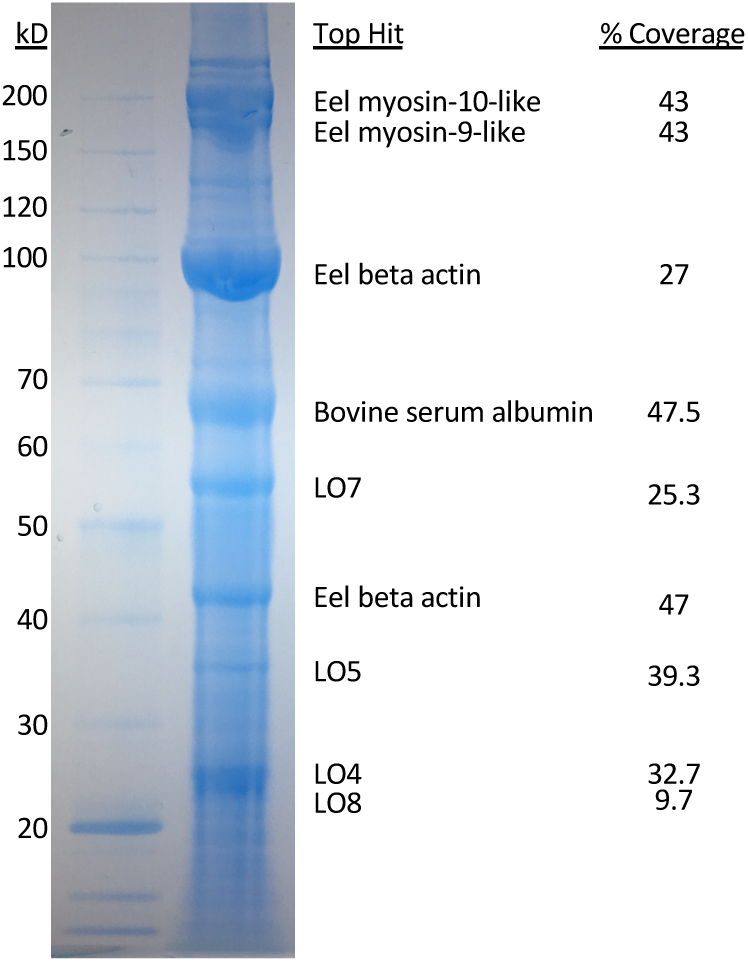
SDS-PAGE analysis of marbled eel adomavirus virions purified from infected EK-1 cells, with bands annotated by mass spectrometric analysis. Clarified lysates of infected cells were ultracentrifuged through an Optiprep gradient. Peak virion-containing fractions were selected then subjected to size exclusion chromatography over agarose resin. The sample was then subjected to TCA precipitation and run on an SDS-PAGE gel. Thirteen gel bands were individually excised, trypsin-digested, and analyzed on a Q Exactive HF Mass Spectrometer. Figure supplement 1: post-translational modifications observed in mass spectrometric results

### Expression of recombinant virion proteins

Adenovirus penton proteins can spontaneously assemble into 12-pentamer subviral particles that may serve as decoy pseudocapsids in vivo (Vragniau, Hubner et al. 2017). Similarly, recombinant polyomavirus and papillomavirus penton proteins can spontaneously assemble into icosahedral virus-like particles (VLPs) that closely resemble native virions. We are not aware of any reports of production of full-size (i.e., hexon+penton) adenovirus VLPs. To investigate the behavior of recombinant adomavirus virion proteins, codon-modified marbled eel adomavirus LO1-LO8 expression plasmids were transfected individually into human 293TT cells (Buck, Pastrana et al. 2004). Optiprep ultracentrifugation was used to separate virus-like particles (VLPs) from smaller solutes. A human papillomavirus type 16 (HPV16) L1/L2 expression plasmid was used as a positive control for VLP formation (Buck, Thompson et al. 2005). Cells transfected with adomavirus LO4, LO5, or LO7 expression constructs each produced particles that migrated into the core fractions of Optiprep gradients, whereas cells transfected with LO1, LO2-3, LO6, or LO8 alone did not show evidence of particle formation (Figure 7). Negative-stain EM analysis showed that the LO4, LO5, and LO7 particles were irregular (Figure 7 Figure supplement 1). In co-transfections of various combinations of LO genes, it was found that inclusion of LO6 inhibited the formation of LO5 and LO7 particles but did not impair the formation of LO4 particles (Figure 8). The fact that over-expression of LO6 can antagonize particle formation supports the bioinformatic prediction that LO6 is a minor virion component that directly interacts with LO5 and LO7.

**Figure 7:**
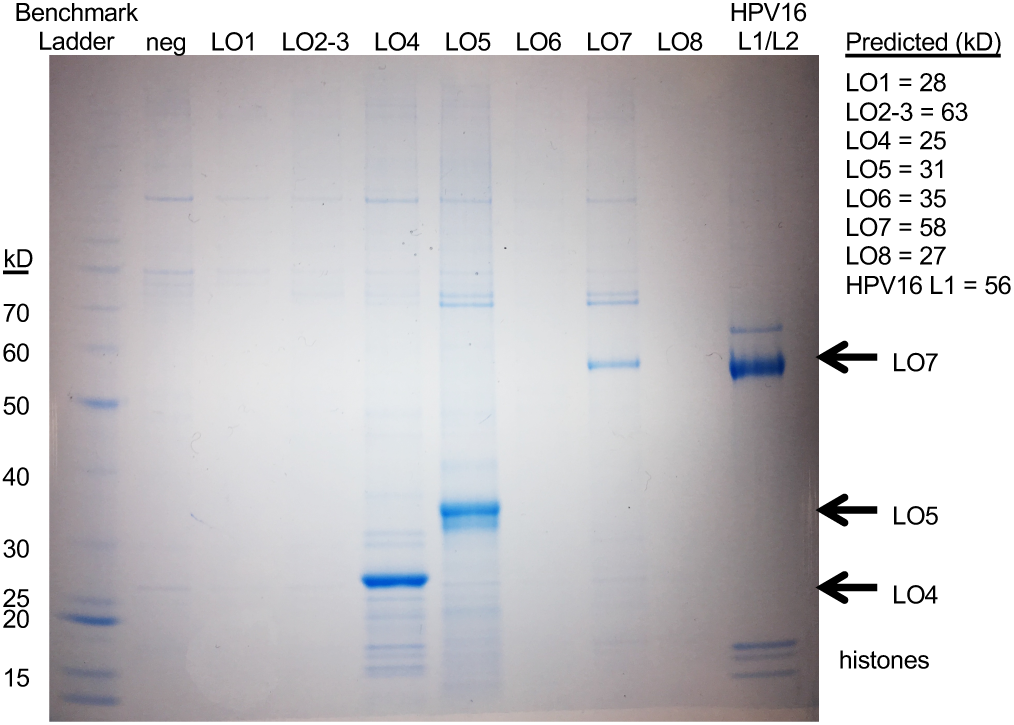
SDS-PAGE analysis VLPs assembled in cells transfected with individual LO expression constructs. 293TT cells were transfected with individual codon modified marbled eel LO expression plasmids indicated at the top of the image. The cells were lysed, subjected to nuclease digestion and a clarifying 5000 x g spin. Soluble material was ultracentrifuged through Optiprep gradients. Core gradient fractions with peak VLP content were subjected to SDS-PAGE analysis. Figure supplement 1: Negative-stain electron microscopy of recombinant particles.

**Figure 8:**
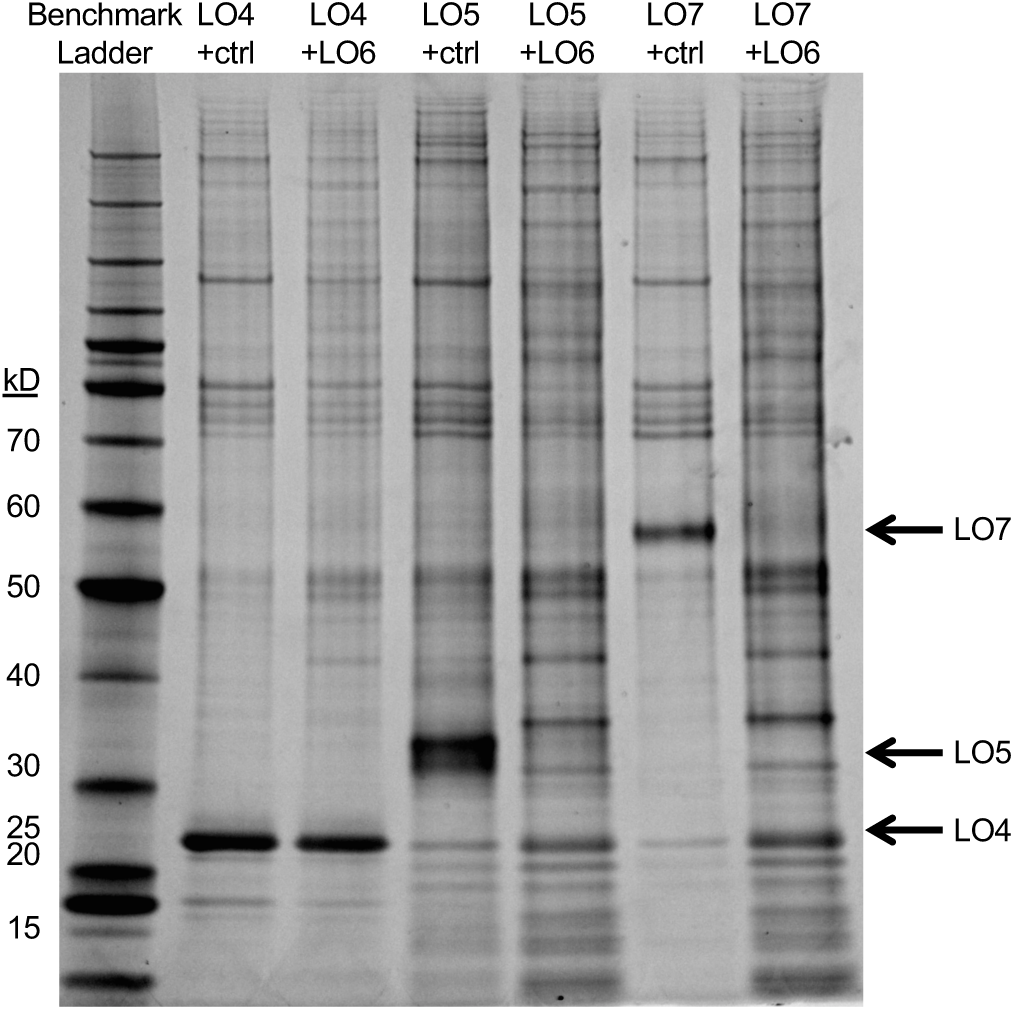
SDS-PAGE gel showing that co-expression of LO6 antagonizes LO5 and LO7 (but not LO4) particle formation. 293TT cells were transfected with the indicated combination of plasmids and subjected to ultracentrifugal separation through Optiprep gradients.

Individually expressed LO5 (penton) and LO7 (hexon) particle preparations showed dsDNA signal in Quant-iT PicoGreen assays (Invitrogen), indicating the presence of nuclease-resistant encapsidated DNA within the purified particles. Optiprep-purified particle preparations from cells co-transfected with LO4, LO5, and LO7 were subjected to an additional round of nuclease digestion with salt-tolerant Benzonase endonuclease (Sigma) followed by agarose gel filtration to remove the nuclease and any residual digested DNA fragments. Nuclease-treated/gel filtered particles typically contained roughly seven nanograms of DNA per microgram of total protein, confirming the presence of nuclease-resistant nonspecific cellular DNA within the particles. The observation is reminiscent of findings for recombinant papillomavirus VLPs (Buck, Thompson et al. 2005).

## Discussion

We have identified a dozen new representatives of the emerging virus family *Adomaviridae*. Four of the new sequences are associated with terrestrial vertebrates, extending the known host range beyond fish. Phylogenetic analyses reveal two adomavirus lineages that appear to have independently co-evolved with host animals. This observation suggests that the two adomavirus lineages both infected the first jawed vertebrates roughly half a billion years ago. Within both adomavirus lineages, there are members associated with commercially important fish species.

Vaccine immunogens comprised of recombinant VLPs have been highly successful in humans (Schiller and Lowy 2015). In particular, vaccines against HPVs have proven remarkably immunogenic, even after a single dose. Our identification of adomavirus virion proteins and demonstration of their ability to assemble into roughly spherical DNA-containing particles should facilitate the development of recombinant subunit vaccines against these viruses, some of which are known to cause severe disease in fish.

The adomavirus virion protein genes, penton (LO5), core (LO6), hexon (LO7), and adenain (LO8), appear to be syntenic homologs of adenovirus and adintovirus virion protein genes. The results also suggest that LO4 may be a homolog of adenovirus pIX, a trimeric coiled coil protein that cements the facets of the adenovirus virion. At a primary sequence level, adomavirus virion proteins more closely resemble adintovirus virion proteins, rather than adenovirus virion proteins (Figure 5). These results tie the *Adomaviridae* into a broad consortium of eukaryotic virus families (Koonin, Krupovic et al. 2015).

In unicellular eukaryotes, non-enveloped midsize (10-50 kb) dsDNA viruses have been shown to have a remarkable degree of genetic modularity (Koonin, Dolja et al. 2015, Yutin, Shevchenko et al. 2015) https://www.biorxiv.org/content/10.1101/697771v3. The pairing of related virion proteins in adenoviruses, adintoviruses, and adomaviruses with entirely different classes of DNA replicase genes thus has ample precedence in non-animal eukaryotes. It will be important to apply emerging higher-throughput search algorithms, such as Mash Screen (Ondov, Starrett et al. 2019) and Cenote-Taker (Tisza, Pastrana et al. 2019), to exhaustively search for each of the overlapping hallmark genes of this virus supergroup in genomic, transcriptomic, and metagenomic surveys, particularly datasets for terrestrial vertebrates.

## Materials and Methods

### Sample Collection and cell culture

A red discus cichlid (*Symphysodon discus*) was purchased at a pet shop in Gainesville, Florida. The fish was moribund and showed erythematous skin lesions. Propagation of the discus adomavirus in cell culture was attempted by overlaying skin tissue homogenates on Grunt Fin (GF) and Epithelioma Papulosum Cyprini cell lines (ATCC). Neither cytopathic effects nor qPCR-based detection of viral replication were observed during two blind passages of 14 days each.

Dr. Chiu-Ming Wen generously provided EK-1 cells (a Japanese eel kidney line) infected with the Taiwanese marbled eel adomavirus (Wen, Chen et al. 2015). The virus was propagated by inoculation of supernatants from the infected culture into uninfected EK-1 cells cultured at room temperature in DMEM with 10 % fetal calf serum. Human embryonic kidney-derived 293TT cells were cultured as previously described (Buck, Pastrana et al. 2004).

### Viral genome sequencing

For the discus adomavirus, total DNA was extracted from a skin lesion and subjected to deep sequencing. Marbled eel adomavirus virions were purified from lysates of infected EK-1 cells using Optiprep gradient ultracentrifugation (Peretti, FitzGerald et al. 2015). DNA extracted from Optiprep gradient fractions was subjected to rolling circle amplification (RCA, TempliPhi, GE Health Sciences). The marbled eel adomavirus RCA products and discus total DNA were prepared with a Nextera XT DNA Sample Prep kit and sequenced using the MiSeq (Illumina) sequencing system with 2 × 250 bp paired-end sequencing reagents. In addition, the marbled eel adomavirus RCA product was digested with *AclI* and *EcoRI* restriction enzymes and the resulting early and late halves of the viral genome were cloned separately into the *AclI* and *EcoRI* restriction sites of pAsylum+. The sequence of the cloned genome was verified by a combination of MiSeq and Sanger sequencing. The clones are available upon request.

For the arowana adomavirus, overlapping PCR primers were designed based on WGS accession numbers LGSE01029406, LGSE01031009, LGSE01028643, LGSE01028176, and LGSE01030049 (Bian, Hu et al. 2016). PCR products were subjected to primer-walking Sanger sequencing.

### Discovery of viral sequences in NCBI databases

Papillomavirus E1 sequences were downloaded from PaVE https://pave.niaid.nih.gov (Van Doorslaer, Li et al. 2017). Polyomavirus LT sequences were downloaded from PyVE https://ccrod.cancer.gov/confluence/display/LCOTF/Polyomavirus (Buck, Van Doorslaer et al. 2016). Parvovirus NS1 proteins and S3H proteins of CRESS viruses and virophage-like viruses were compiled from multiple databases, including RefSeq, WGS, and TSA, using TBLASTN searches. Adenovirus, virophage, and bacteriophage PolB sequences were downloaded from GenBank nr using DELTA-BLAST searches (Boratyn, Schaffer et al. 2012) with Alpha or Beta adintovirus PolB proteins as bait.

SRA datasets for fish, amphibians, and reptiles were searched using DIAMOND (Buchfink, Xie et al. 2015) or NCBI SRA Toolkit (http://www.ncbi.nlm.nih.gov/books/NBK158900/) in TBLASTN mode using adomavirus protein sequences as the subject database or query, respectively. Reads with similarity to the baits were collected and subjected to BLASTX searches against a custom library of viral proteins representing adomaviruses and other small DNA tumor viruses. SRA datasets of interest were subjected to de novo assembly using the SPAdes suite (Bankevich, Nurk et al. 2012, Nurk, Meleshko et al. 2017) or Megahit (Li, Liu et al. 2015, Li, Luo et al. 2016). Contigs encoding virus-like proteins were identified by TBLASTN searches against adomavirus protein sequences using Bowtie (Langmead and Salzberg 2012). The candidate contigs were validated using the CLC Genomics Workbench 12 align to reference function.

Predicted protein sequences were automatically extracted from contigs of interest using getorf (http://bioinfo.nhri.org.tw/cgi-bin/emboss/getorf)(Rice, Longden et al. 2000). Sequences were clustered using EFI-EST (https://efi.igb.illinois.edu/efi-est/)(Gerlt, Bouvier et al. 2015, Zallot, Oberg et al. 2018) and displayed using Cytoscape v3.7.1 (Shannon, Markiel et al. 2003). Multiple sequence alignments were constructed using MAFFT (https://toolkit.tuebingen.mpg.de/#/tools/mafft)(Kuraku, Zmasek et al. 2013, Katoh, Rozewicki et al. 2019). Individual or aligned protein sequences were subjected to HHpred searches (https://toolkit.tuebingen.mpg.de/#/tools/hhpred)(Hildebrand, Remmert et al. 2009, Meier and Soding 2015, Zimmermann, Stephens et al. 2017) against PDB, Pfam-A, NCBI Conserved Domains, and PRK databases.

Contigs were annotated using Cenote-Taker (Tisza, Pastrana et al. 2019) with an iteratively refined library of conserved adintovirus protein sequences. Maps were drawn using MacVector 17. Phylogenetic analyses were performed using Phylogeny.fr with default settings (Dereeper, Guignon et al. 2008).

### Marbled eel adomavirus transcript analysis, late ORF expression, and virion purification

RNAseq reads reported by Wen et al (Wen, Chen et al. 2015) were aligned to the marbled eel adomavirus genome using HISAT2 version 2.0.5 (Kim, Langmead et al. 2015) with the following options: “--rna-strandness FR --dta --no-mixed --no-discordant”. Integrated Genome Viewer (IGV) version 2.4.9 (Robinson, Thorvaldsdottir et al. 2017) was used to determine splice junctions and their depth of coverage. Additional validation was performed by visual inspection using CLC Genomics Workbench 12.

Codon-modified expression constructs encoding the marbled eel adomavirus LO1-LO8 proteins were designed according to a modified version of a previously reported algorithm (https://github.com/BUCK-LCO-NCI/Codmod_as_different_as_possible)(Pastrana, Buck et al. 2004). 293TT cells were transfected with LO expression constructs for roughly 48 hours. Cells were lysed in a small volume of PBS with 0.5% Triton X-100 or Brij-58 and Benzonase Dnase/Rnase (Sigma)(Buck and Thompson 2007). After one hour of maturation at neutral pH, the lysate was clarified at 5000 x g for 10 min. The clarified lysate was loaded onto a 15-27-33-39-46% Optiprep gradient in PBS with 0.8 M NaCl. Gradient fractions were collected by bottom puncture of the tube and screened by PicoGreen dsDNA stain (Invitrogen), BCA, or SDS-PAGE analysis. Electron microscopic analysis was performed by spotting several microliters of Optiprep fraction material (or, in some instances, particles exchanged out of Optiprep using agarose gel filtration) onto carbon film copper grids, followed by staining with 0.5% uranyl acetate.

### Mass Spectrometry

Optiprep-purified marbled eel adomavirus virions were precipitated with trichloroacetic acid. A 1 ml sample was treated with 100 µl of 0.15% deoxycholic acid and incubated at room temperature for 10 minutes. 100 µl of 100% TCA was then added and the sample was vortexed and incubated on ice for 30 minutes. Following the incubation, the sample was centrifuged at 10,000 x g for 10 minutes at 4°C. The supernatant was removed, and the remaining pellet was washed with ice-cold acetone to remove residual TCA. The protein pellet was solublized with NuPAGE Sample Buffer + 5% BME (Sigma) and run on a 10-12% Bis-Tris MOPS gel (Thermo). The protein bands were visualized using InstantBlue (Expedeon). Thirteen gel bands were individually excised and placed into 1.5 ml Eppendorf tubes. The gel bands were sent to the National Cancer Institute in Fredrick, Maryland where they were de-stained, digested with trypsin, and processed on a Thermo Fisher Q Exactive HF Mass Spectrometer. Thermo Proteome Discoverer 2.2 software was used for initial protein identification. The uninterpreted mass spectral data were also searched against *Anguilla* proteins (Swiss-Prot and TrEMBL database containing 105,268 proteins), Bos taurus proteins (Swiss-Prot and TrEMBL database containing 48,288 proteins), a common contaminants database (cRAPome), and translated marbled eel adomavirus ORFs. Further analysis was conducted using Protein Metrics Biopharma software to identify modifications missed in initial analyses.

## Supporting information

Fig2Supp1 Accession Numbers

Fig2Supp2 examples of annotation

Fig2Supp3 Nucleotide Maps

Fig2Supp4 Adoma Protein Compilation

Fig2Supp5 Adoma splicing

Fig4Supp1 S3H Network File

Fig4Supp2 LO8 Tree

Fig5Supp1 Minor Virion Protein Network Files

Fig6Supp1 Mass Spec PTMs

Fig7Supp1 VLP Electron Microscopy

## Ethics Statement

All animal tissue samples were received as diagnostic specimens collected for pathogen testing and disease investigation purposes.

## Data Availability

GenBank accession numbers for sequences deposited in association with this study are BK010891 BK010892 BK011012 BK011013 BK011014 BK011015 BK011016 BK011017 BK011018 BK011019 BK011020 BK011021 BK012039 BK012040 BK012041 MF946549 MF946550 MH282863.

## Acknowledgments

The authors are indebted to Eugene Koonin and Natalya Yutin for their generous guidance and for the spirited discussions that inspired us to pursue this study. We are particularly grateful to them for sharing their observation that adomaviruses encode a recognizable adenain homolog and their discovery of adomavirus sequences in the arowana WGS datasets. The authors are also grateful to Lisa Jenkins for her extensive technical guidance on analyzing mass spectrometric data. We thank Karl Münger for useful discussions, including advice about oncogene sequence motifs.

