## Supplementary material for "Identification of Adomavirus Virion Proteins": Fig2Supp2 examples of annotation

The predicted SET protein of Discus adomavirus was submitted for searches of PDB_mmCIV70, COG_KOG, Pfam-A, and NCBI_CD databases in HHPred https://toolkit.tuebingen.mpg.de/tools/hhpred. The figure shows a screen shot of the results.

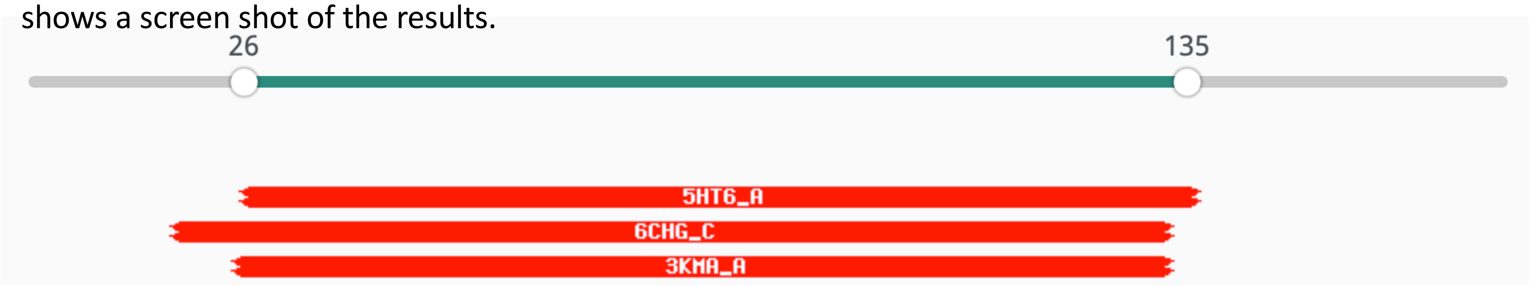


### Discus SET (HHpred)

3. **3KMA_A A612L protein; SET domain, VIRAL PROTEIN; HET: GOL, SO4; 1.6A {Paramecium bursaria Chlorella virus 1} SCOP: b.85.7.2; Related PDB entries: 3KMJ_A 3KMT_B 3KMT_A 1N3J_B 2G46_B 3KMT_C 1N3J_A 2G46_A 3KMA_B**

Probability: 97.47%, E-value: 0.000055, Score: 47.59, Aligned cols: 98, Identities: 16%, Similarity: 0.17,

```
Q ss_pred               CCCEEEEeCCCCccEEEEecCcCCCCEEEEccCcccccccceEeEEEEEEEecCCCCCcccCCCcccCCCCccceeeecCCC
Q Q_1452582     25      DPPIIVITHPKKGEIVIAKRNISKGDIVFCWPGSVTTGLGVRLQNVTFEVPYPRPPFFTQAPAKENTEPLKCSYFTEDSE   104 (172)
Q Consensus     25      dppiivithpkkgeiviakrniskgdivfcwpgsvttglgvrlqnvtfevpyprppfftqapakenteplkcsyftedse   104 (172)
                        .+++.+-..+.+|.=+.|.+.|.+|+++..++|.+....-..-+.-.+-..+...          ....+....|-..+.
T Consensus     3       ~~~~~v~~~~~~g~gl~a~~~i~~g~~i~~~~g~~~~~~~~~~~~~~~~~~~~~~----------~~~~~~~~~~~nhs~   72 (119)
T 3KMA_A        3       NDRVIVKKSPLGGYGVFARKSFEKGELVEECLCIVRHNDDWGTALEDYLFSRKNM----------SAMALGFGAIFNHSK   72 (119)
T ss_dssp               CSSEEEEECTTSSEEEEESSCBCTTCEEEEEEEEEEEGGGCCSTTTTTCEEETTE----------EEEETBTTTTCEECS
T ss_pred               CCcEEEEeCCCCceEEEECCccCCCCEEEEEEEEEEEeCccccchhHHheeecCCc----------eeEecCeeeeeecCC


Q ss_pred               CCCceEEE-----EEEEEEeCCCCCCceEEEee
Q Q_1452582     105     GPNGTFSI-----GLFFKATTDIEPGRQVFYDF   132 (172)
Q Consensus     105     gpngtfsi-----glffkattdiepgrqvfydf   132 (172)
                        .||-.+..     .+.+.|..+|++|.+++.|+
T Consensus     73      ~pn~~~~~~~~~~~~~~~~~a~~~i~~g~el~~~Y   105 (119)
T 3KMA_A        73      DPNARHELTAGLKRMRIFTIKPIAIGEEITISY   105 (119)
T ss_dssp               SCSEEEEECTTSCEEEEEEESSCBCTTCEEEECC
T ss_pred               CCceEEEEeCCCceEEEEEEECCccCCCCeeEEec
```

### Discus SET (NCBI BLASTP)

| | Description | Max Score | Total Score | Query Cover | E value | Per. Ident | Accession |
|---|---|---|---|---|---|---|---|
| ☑ | SET domain-containing protein [Hyphomicrobium sp. 99] | 44.3 | 44.3 | 70% | 0.050 | 26.77% | WP_045835396.1 |
| ☑ | SET domain-containing protein-lysine N-methyltransferase [Hyphomicrobium sp.] | 43.5 | 43.5 | 67% | 0.10 | 25.81% | RUP09790.1 |
| ☑ | hypothetical protein LBMAG06_11890 [Actinobacteria bacterium] | 42.7 | 42.7 | 66% | 0.14 | 26.89% | GDX19060.1 |
| ☑ | SET domain-containing protein [Hyphomicrobium denitrificans] | 42.0 | 42.0 | 70% | 0.32 | 27.27% | WP_041319975.1 |
| ☑ | MULTISPECIES: SET domain-containing protein [unclassified Hyphomicrobium] | 42.0 | 42.0 | 70% | 0.36 | 25.58% | WP_013948827.1 |
| ☑ | hypothetical protein ABR56_06210 [Acidimicrobium sp. BACL27 MAG-120823-bin4] | 40.8 | 40.8 | 63% | 0.65 | 27.19% | KRO47834.1 |
| ☑ | TPA: SET domain-containing protein-lysine N-methyltransferase [Acidimicrobium sp.] | 40.4 | 40.4 | 63% | 1.0 | 27.19% | HBZ62264.1 |
| ☑ | SET domain-containing protein-lysine N-methyltransferase [Hyphomicrobium sp.] | 39.7 | 39.7 | 67% | 1.9 | 25.81% | RUO98239.1 |
| ☑ | hypothetical protein AK812_SmicGene42166 [Symbiodinium microadriaticum] | 40.4 | 40.4 | 30% | 2.0 | 38.46% | OLP77740.1 |
| ☑ | SET domain-containing protein [Hyphomicrobium facile] | 38.5 | 38.5 | 67% | 5.1 | 26.61% | WP_092867925.1 |
| ☑ | related to Allantoinase [Hanseniaspora guilliermondii] | 38.5 | 38.5 | 46% | 7.4 | 28.75% | SGZ39326.1 |
| ☑ | Allantoinase [Hanseniaspora uvarum] | 38.5 | 38.5 | 20% | 8.7 | 42.86% | OEJ84692.1 |
| ☑ | nuclear protein SET [Hyphomicrobium denitrificans 1NES1] | 37.4 | 37.4 | 56% | 9.7 | 26.92% | AGK57099.1 |

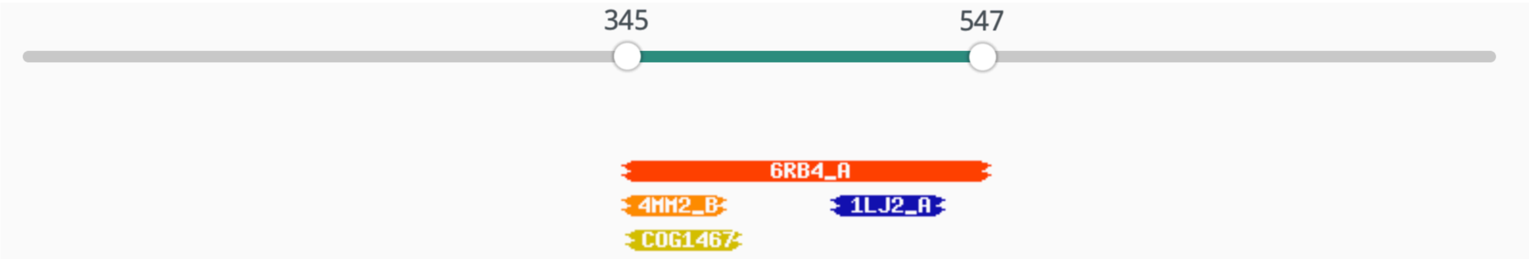


### Discus EO3 (AEP)

1. **6RB4_A DNA primase small subunit (E.C.2.7.7.-); Primase, DNA-dependent RNA polymerase, ATP; HET: EDO; 1.5A {Homo sapiens}; Related PDB entries: 4MHQ_A 4BPX_C 4BPX_A 4BPU_C 4BPU_A 4BPW_C 4BPW_A 5EXR_E 5EXR_A 4RR2_C 4RR2_A 6R4S_E 6R5D_E 6R4S_A 6R5D_A 6R4T_D 6R5E_E 6R4T_A 6R5E_A 6R4U_E 6R4U_A 4LIK_A 4LIL_A**

   Probability: 94.31%, E-value: 0.12, Score: 55.1, Aligned cols: 192, Identities: 18%, Similarity: 0.3,

```
Q ss_pred             hhhccCCcchHHHHHHHHHHHHHHHHHHHHHHHHHHhheeEEecCCceEEEEEcCccccchhhcccccchhhHhcCh--
Q Q_7558216     345  RACCRDTNTMCTYCCQVLTFAADLLLAFLVAMYAAQHVQVYYDGNCGLFVVCHDKHLHMTKTEMCTMQKNWETVLTNP--    422 (839)
Q Consensus     345  raccrdtntmctyccqvltfaadlllaflvamyaaqhvqvyydgncglfvvchdkhlhmtktemctmqknwetvltnp--    422 (839)
                       |.||.+. ..|..|-.....|..++-.+|-.-+.-.++.++|.|+-|.-+...|+....-..+.-.---++-++....
T Consensus     122  R~Cc~~~-~iC~~Cw~~~~~A~kll~~~L~~dfGf~~~~~vFSG~RG~H~~V~D~~ar~L~~~~R~aIv~Yl~~~~~~~~    200 (410)
T 6RB4_A        122  RRCCSSA-DICPKCWTLMTMAIRIIDRALKEDFGFKHRLWVYSGRRGVHCWVCDESVRKLSSAVRSGIVEYLSLVKGGQD    200 (410)
T ss_dssp             CSSSCTT-CBCTTTHHHHHHHHHHHHHCCCCCTCCCCEEEEECSSSEEEEEECCHHHHTCCHHHHHHHHHHHCCCCCCTT
T ss_pred             ccccCCc-ccCHHHHHHHHHHHHHHHHHHHHHHhhCCCceEEEEecCCceEEEECCHHHhcCCHHHHHHHHHHHheeeCCCc


Q ss_pred             --------hhCHHHHHHHhHHHHhccEEEEEeeecccCCcchHHHH-------HHHHHHHhCCCCHHHHHHHHHHHHHHH
Q Q_7558216     423  --------SAYPEFIRAADQAVRRSAVVIASVKFSKRNRPYAASAI-------MLQEFLRSMDLPQEREKRVFEDMRRFI    487 (839)
Q Consensus     423  --------saypefiraadqavrrsavviasvkfskrnrpyaasai-------mlqeflrsmdlpqerekrvfedmrrfi    487 (839)
                               ......++.|-...+....-+.-..-.--..+-....+       ..+++...........| ++.+..++
T Consensus     201  ~~~~~~~~~~~hp~~~~a~~~~~~~~f~~~~~~~~~q~~~~~~~~~~~~~~~~l~~~~~~~~~~~~~l~~~~~~~~~~~~r-w~~l~~~~    279 (410)
T 6RB4_A        201  VKKKVHLSEKIHPFIRKSINIIKKYFEEYALVNQDILENKESWDKILALVPETIHDELQQSFQKSHNSLQR-WEHLKKVA    279 (410)
T ss_dssp             CSCCCCCCSSCCHHHHHHHHHHHHHHCCCCCCCSCTTSSHHHHHHHHTTSCGGGHHHHHHHHHHSCSHHHH-HHHHHHHH
T ss_pred             ccceecccccCHHHHHHHHHHHHHHHHHhhcccccccChhhHHHHHhhCHHHHHHHHHHHHHHHhCCCHHHH-HHHHHHHH


Q ss_pred             HHhCCCCcchhhhHHHHHHhhhcchHHHHHHHHHHhcceeehhc---cccCCcccEEEEeCCCch
Q Q_7558216     488  RKYEPNDADMRDTALYFVSLFSGYEFAVHMLKALRPCVHVNT---LHKGFPVPFMVHEGTGNL    547 (839)
Q Consensus     488  rkyepndadmrdtalyfvslfsgyefavhmlkalrpcvhvnt---lhkgfpvpfmvhegtgnl    547 (839)
                       .++..............          ..-.+|+-+.|-+-+|.   ++.=...||-||.+||..
T Consensus     280  ~~~~~~~~~~~~~~~~~~---------~~~ivl~~~yPrLD~~VS~~~nHLLK~PfcVHpKTG~v    333 (410)
T 6RB4_A        280  SRYQNNIKNDKYGPWL---------EWEIMLQYCFPRLDINVSKGINHLLKSPFSVHPKTGRI    333 (410)
T ss_dssp             HTCCSSSSSCCCCSHH---------HHHHHHHHHSCCCCHHHHHCTTCCEECTTCBCTTTCBB
T ss_pred             HHHhccccCccchHH---------HHHHHHHHhcCccCccccccccHhhcCCCccCCCCCCcE
```

| | BLASTP Description | Max Score | Total Score | Query Cover | E value | Per. Ident | Accession |
|---|---|---|---|---|---|---|---|
| ☑ | AGAP005563-PA-like protein [Anopheles sinensis] | 44.7 | 44.7 | 11% | 1.6 | 29.70% | KFB50766.1 |
| ☑ | hypothetical protein HCUR_01017 [Holospora curviuscula] | 38.9 | 38.9 | 5% | 3.7 | 39.58% | PPE03479.1 |

### Multiple alignment of LO5

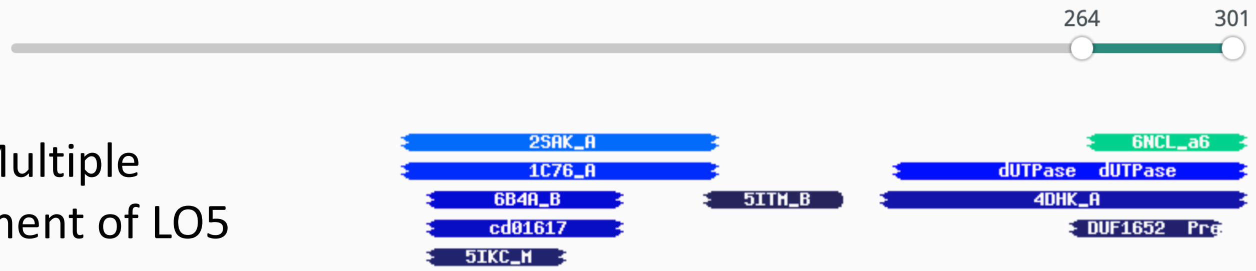


#### Hitlist

Show 25 Entries Search:

| Nr | Hit | Name | Probability | E-value | SS | Cols | Target Length |
|---|---|---|---|---|---|---|---|
| 1 | 6NCL_a6 | PBCV-1 capsid; tape-measure protein, minor capsid proteins; 3.5A {Paramecium bursaria Chlorella virus 1} | 61.68 | 37 | 4.6 | 38 | 170 |
| 2 | 2SAK_A | STAPHYLOKINASE, 2-AMINO-2-HYDROXYMETHYL-PROPANE-1,3-DIOL; PLASMINOGEN ACTIVATION, FIBRINOLYSIS, STAPHYLOKINASE, HYDROLASE; 1.8A {Staphylococcus aureus} SCOP: d.15.5.1 | 49.09 | 110 | 5 | 72 | 121 |
| 3 | 1C76_A | STAPHYLOKINASE; BETA-GRASP FAMILY, HYDROLASE; 2.25A {Staphylococcus aureus} SCOP: d.15.5.1; Related PDB entries: 1C77_B 1C77_A 1C78_B 1C78_A 1C79_B 1C79_A 1SSN_A 1BUI_C | 44.13 | 130 | 5 | 72 | 136 |
| 4 | PF00692.19 | ; dUTPase ; dUTPase | 39.99 | 370 | 6.6 | 78 | 128 |

Template alignment | Template 3D Structure | PDBe

1. **6NCL_a6 PBCV-1 capsid; tape-measure protein, minor capsid proteins; 3.5A {Paramecium bursaria Chlorella virus 1}**
   Probability: 61.68%, E-value: 37, Score: 32.6, Aligned cols: 38, Identities: 24%, Similarity: 0.33,

```
Q ss_pred                ccCCccCEEEEEEEccCCCcceeeecCee-EEEEEEeCC
Q LO5_Carp_Adoma    264  HERPTIPELTFRLRDRANRPLSFLSGLPC-AIIELMHLD   301 (301)
Q Consensus         264  ~~~p~l~~Ltf~L~Dr~~Rpl~FvsGvPs-ati~i~~L~   301 (301)
                         .+..+|.+|+++++|..|..+.|-.|--- -+||+.+++
T Consensus         111  ~PI~kLDrLtI~~rd~nG~~i~f~~~~~N~sF~lrF~c~~   149 (170)
T 6NCL_a6           111  NPVATMDKLNIKLKDANGNVLTIPAGNEHSFMIQLTCGD   149 (170)
T ss_dssp                 EEEEEEEEEEEEEECTTSCBCCCCTTCCCEEEEEEEECT
T ss_pred                 CccccccEEEEEEECCCCCEeeccCCCCeeEEEEEEeCC
```

### PSI-BLAST of LO5 MSA

| | | |
|---|---|---|
| GBM63801.1 hypothetical protein AVEN_262691_1 [Araneus ventricosus] | 60.8 | 5e-07 |
| GBN35576.1 hypothetical protein AVEN_162585_1 [Araneus ventricosus] | 57.8 | 5e-06 |
| GBM20513.1 hypothetical protein AVEN_257516_1, partial [Araneus v... | 54.3 | 6e-05 |
| EFA12278.1 hypothetical protein TcasGA2_TC005263 [Tribolium casta... | 55.1 | 1e-04 |
| EFA11812.1 hypothetical protein TcasGA2_TC008591 [Tribolium casta... | 52.4 | 8e-04 |
| GBN06273.1 hypothetical protein AVEN_265340_1 [Araneus ventricosus] | 49.7 | 0.006 |
| WP_143457370.1 hypothetical protein, partial [Klebsiella pneumoniae] | 45.8 | 0.016 |
| GBN69182.1 hypothetical protein AVEN_76811_1 [Araneus ventricosus] | 47.4 | 0.022 |
| EFA09107.1 hypothetical protein TcasGA2_TC015482 [Tribolium casta... | 47.0 | 0.049 |
| GBN36035.1 Uncharacterized protein F54H12.2, partial [Araneus ven... | 47.0 | 0.060 |
| GBM99268.1 hypothetical protein AVEN_272123_1 [Araneus ventricosus] | 45.4 | 0.10 |

Barramundi LO6

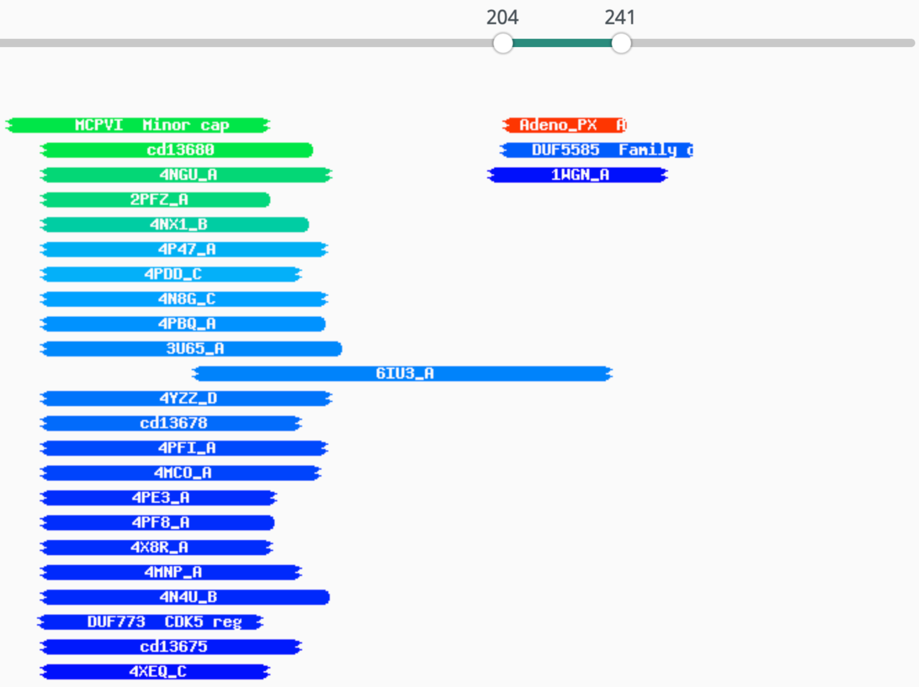


#### Hitlist

Show 25 Entries Search:

| Nr | Hit | Name | Probability | E-value | SS | Cols | Target Length |
|---|---|---|---|---|---|---|---|
| 1 | PF05829.12 | ; Adeno_PX ; Adenovirus late L2 mu core protein (Protein X) | 95.32 | 0.0069 | 2.8 | 38 | 42 |
| 2 | PF02993.14 | ; MCPVI ; Minor capsid protein VI | 65.71 | 45 | 6.1 | 65 | 221 |
| 3 | cd13680 | PBP2_TRAP_SBP_like_4; Uncharacterized substrate-binding protein of the Tripartite ATP-independent Periplasmic transporter family. | 65.65 | 110 | 7.7 | 84 | 300 |
| 4 | 4NGU_A | TRAP dicarboxylate transporter, DctP subunit; TRAP periplasmic solute binding family; HET: DAL, MSE; 2.5A {Desulfovibrio alaskensis} SCOP: l.1.1.1, c.94.1.0 | 62.98 | 150 | 8.3 | 89 | 319 |

Template alignment | CDD

1. **PF05829.12 ; Adeno_PX ; Adenovirus late L2 mu core protein (Protein X)**

Probability: 95.32%, E-value: 0.0069, Score: 45.78, Aligned cols: 38, Identities: 26%, Similarity: 0.441,

```
Q ss_pred                hCCHHHhcccHHHHHHHHHHHHHHhccchHHHHHHHHHH
Q Q_8406900     204    DMPKSDRGGILPFIIPAIATVLGATIPQIPKFIQLARD    241 (333)
Q Consensus     204    ~M~k~eRgg~ip~lip~l~~~l~~~~~~ipkfiqla~d    241 (333)
                         ....+-||||+|+|||+|+.+||+.-+--.-.||....
T Consensus     5      ~~~~~lrGGflpaLiPiiaaaIgAiPgIAs~ai~a~n~    42 (42)
T PF05829.12    5      SYNKELRGGFLPLLVPIIAAAISAAPGIAGAVLAAKNH    42 (42)
T ss_pred                ccccccccccHHHHHHHHHHHHHHhhhHHHHHHHHHhcC
```

Template alignment | CDD

2. **PF02993.14 ; MCPVI ; Minor capsid protein VI**

Probability: 65.71%, E-value: 45, Score: 33.44, Aligned cols: 65, Identities: 23%, Similarity: 0.386,

```
Q ss_pred                ccCCCcHHHHHHHHHHHHHhcHHHHHHhHHHHHHHHHHHH--HHCHHHHHHHHhcCCchHhHhhHhchHHHHHHHHH
Q Q_8406900     50     RGGRFNFKELALKAWKNLKASFFPIVRQKGKEFLQTAADHAV--KKGPQYYEALKKDGVTGVRDMFLDNVSTVATDFVRK    127 (333)
Q Consensus     50     RGGg~n~~~lA~kA~~~LKan~vPli~r~~k~~~q~aaehAv--knG~qyy~AfKk~GV~Gv~d~vldnlp~v~gd~~r~    127 (333)
                         |||-|||..|-.+-|++||         +.++|++..+..|.  +-|.+.-+-||+.|       |.+|+-..+++-|..
T Consensus     24     ~GGAfnWgslgs~~wSgLk----------~~Gs~ikn~g~k~~nS~t~q~lkq~lkDsg-------v~ekvv~gag~gi~g    87 (221)
T PF02993.14    24     RGGAINWGNLGSRITSALN---------STGRWLYNTGNRFVHSNTFNQIKQGIQDSG-------VIRNVANLAGETLGA    87 (221)
T ss_pred                CCCccccchhcchHhHHHHH---------HhhHhHHHHhhhchhccHHHHHHHHHHHhhhc-------HHHHHHHHHHHHHHHH
```

Resubmit Section

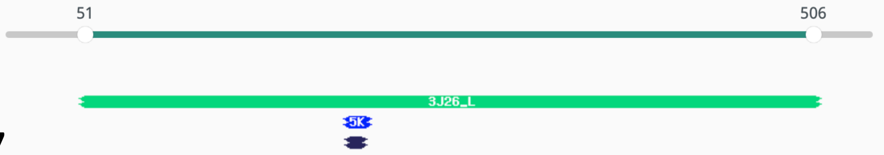


### Grenadier LO7

#### Hitlist

Show 25 Entries Search:

| Nr | Hit | Name | Probability | E-value | SS | Cols | Target Length |
|---|---|---|---|---|---|---|---|
| 1 | 3J26_L | capsid protein V20, Minor virion; double jelly-roll, single jelly-roll, VIRUS; 3.5A {Sputnik virophage}; Related PDB entries: 3J26_I 3J26_A 3J26_M 3J26_D 3J26_J 3J26_C 3J26_H 3J26_E 3J26_B 3J26_G 3J26_K 3J26_F | 62.55 | 640 | 17.8 | 413 | 508 |
| 2 | 5KUH_B | Glutamate receptor ionotropic, kainate 2; GluK2EM with LY466195, SIGNALING PROTEIN; HET: LY5; 11.6A {Rattus norvegicus}; Related PDB entries: 5KUH_D 5KUH_C 5KUH_A | 43.16 | 26 | 0.5 | 16 | 757 |
| 3 | PF16926.5 | ; HisKA_4TM ; Archaeal 4TM region of histidine kinase | 23.74 | 49 | -1.1 | 12 | 165 |

#### Alignments

Template alignment | Template 3D Structure | PDBe

1. **3J26_L capsid protein V20, Minor virion; double jelly-roll, single jelly-roll, VIRUS; 3.5A {Sputnik virophage}; Related PDB entries: 3J26_I 3J26_A 3J26_M 3J26_D 3J26_J 3J26_C 3J26_H 3J26_E 3J26_B 3J26_G 3J26_K 3J26_F**

Probability: 62.55%, E-value: 640, Score: 30.9, Aligned cols: 413, Identities: 12%, Similarity: 0.06,

```
Q ss_pred             eehhcCccc--eEEEEeeCCCcce------eEeeeEEEEEEeeEEEEeCCCcceeeeEecCCCCC-cceeccCCCCcCCCC
Q Q_GrenLO7    51    QLNSRGTS--DVSFHLQCPRDEA------ILQGSIQWVCEGRVAVITAATGAEVVWQTAGQPGN-GAVQFNCGPNAVNNN   121 (543)
Q Consensus    51    qlN~~gt~--e~~i~lqC~~DEA------IL~GS~~fva~G~V~~~~~~~~~~~~~~WQs~~~~~~~~~~~~~~~~~~~~~N   121 (543)
                      +.|.-+.+  .+.+.++||..--      .+.+.+.|..+|++. ...-|..+-+|-- ++-+. -..++-..-++.=+|
T Consensus    39    ~~na~s~S~ss~tf~i~~PSe~~v~dR~i~~~~~v~~t~t~t~~~~~pig~~l~~~g~~al~afPl~s~~~t~~~tiNn   116 (508)
T 3J26_L      39    PFPSTSFSSNQFNFICNPPSAQTVLDRLVFIQVPYDITFTANPS-HAGITENLLQPGR-DAFRAFPISSITNTLNATING   116 (508)
T ss_dssp            EEECSSCCSSEEEEECCCSSSSCEECSCCEEEEEEEEEEECCSS-SCCCCSCSSCTTT-EEECSSHHHHHEEEEEEEETT
T ss_pred             eeccccccCceeEEEEeCCccceeEeeeEEEeecEEEEEEecCC-CCCCcccccccch-HhccccHHHHHhheeeeeCC

Q ss_pred             cCCCCcCCCceeecccceEEEEEecCChHH-HHHhhhhcccccCcchHHHHHHHhcCCCCCCcccccCCcCcccccceeece
Q Q_GrenLO7    122   AAAPAVNIPPHVVSSQFTQIDIETGDTPSL-NRHLAQVSASECGAGALSAVNAYLNETANPGYYALNSTTVAGLVHRHGL   200 (543)
Q Consensus    122   ~~~~~vpiPpfv~NsQFqqID~e~G~t~S~~n~~L~~~rsS~~g~gal~~~N~Y~N~p~~~G~yhenst~~~~~~~~~~~~~~   200 (543)
                        ..+.|++-=+--|+.    ++.+.--+ .+++...-+=..--..-+-+---.|.|.....-..+....--+..+...
T Consensus    117   -----vsin~~di~~~ll----r~~d~~~~~~~~~~~s~tPSm~D~Y~sY~da~~~~nNPl~~~~~~s~d~~~pRGsf~~~~   189 (508)
T 3J26_L      117   ---FPVNIELAQIIHALS----RYHTPLKVKNGWMSMQPSFEDNYQSYRDADGANNNPLGVFTSAAGLSELPRGSYTMNV   189 (508)
T ss_dssp            ---EEEEECCTTTTHHHH----TTTCCHHHHTTGGGGSCCCCCCSSSHHHHCSCSSCTTCCSTTSCTTSCCCGGGSCCEE
T ss_pred             ---ccceecHHHHHHHHH----HhcCccchhcCcccCCCCChhhcchhHHHHhcccCCCCcccccCCccCCCCeeEEEEE

Q ss_pred             eecCCCCceeeeeeehhccCCCcceeeeecCCCcccceecCCCCCcEEEEEeCCchhheeeeeCCC----CceEEEEEEe
Q Q_GrenLO7    201   ALNRSTDVFNCVERVHHQINLPFSMLCSWDYDDNHTRKQYPRGMSLRIRCSTRNWQSRSRLATSQD----GLHYAHIVWT   276 (543)
Q Consensus    201   aV~~~~d~f~~~Erv~HqLN~P~N~L~swd~dD~etrk~lPrGm~lrir~~tr~~~~R~~l~~~~d----~~~~~~~~~i~f~   276 (543)
                      -++..++.      ..--.+.-|+ +|.-..|.|.+.- -+.-==.+.+.+.-..-+.|+.=.++.+    +.-++..-..
T Consensus    190   ~~~t~t~~~~~~~~It~~~~Epl~~lSPfv~~~~~~~~~~G~~gl~t~~f~~~l~~~~~rm~s~s~~~~~~~~~~ati~~~~~   262 (508)
T 3J26_L      190   VTNTTTTA-----RITGVLYEQV-FLPPFLWDGEQAG-GLANLTSLTFNWVLNNNLARIWSHSDITNDVSGNSTIGSMNI   262 (508)
T ss_dssp            EEEETTEE-----EEEEEEEECC-CSTTSCCSSSCCC-CCCCCSCCEEEEEECSCCCTTEEECGGGTCSSSCCSCCCEEE
T ss_pred             EecCCceE-----EEEEEEecce-eeCceeccCCCcC-ccccceeEEEEEcCCchHHhhccccCCCCCCCceecccch

Q ss_pred             EE-----EEEEEEEECCCCccccCCCceeeeeeeeeEeeEec---CCCCCCeEEEEcccCCchhcccEEEEEEcccccCC
Q Q_GrenLO7    277   KV-----TIKYTTATVPDGAHRQAEPLETVFATLDMGIRMYQ---PEANNPHMTIHITTSPTEFIPQWVIVWLAPADCFD   348 (543)
Q Consensus    277   ~~~~~~~~~v~ytti~vpd~~hR~~Ep~~t~yptmD~gvr~~~~~~~l~agq~s~~ipIT~sd~d~vPQ~lv~fvA~~d~F~   348 (543)
                      .|      -|.|-   =|....  .-|.+.+||-+++.-..-+   +.+.+.+..+---.-..+-||-+|++|+-..+.|-
T Consensus    263   ~~~~~~sllv~f~----P~~~~~~~~i~~~~~v~P~~e~~ry~T~~s~~~~p~as~~~~S~~IqLn~VPd~l~I~vr~~~~~~~   337 (508)
T 3J26_L      263   SFQQPSMYLGFV---TPRLNI--PIPPRITYPYFKLSRYTTQFQNTLAPNASSTFKSNVVQLDSIPRKLYLFVKQSDNVI   337 (508)
T ss_dssp            EECCCEEEECEE---CCCTTS--CCCSSEEEEEEEEEEEEEECCCCBCSSCBCCCCCCCEECSSCCSEEEEEEEEECSHHH
T ss_pred             hcCCceeeeeec---CcccCC--CCCCcceeecchhcceecCcCCCCCCCcceeeeceeecccCCcEEEEEEccccccc

Q ss_pred             hhhcccccccceecccchhhEEEEecCCCCCcchhhheeccceeChhHHHHHHHHHHHHHHHHhcCCCCCCCCCcCCHHHHhH
Q Q_GrenLO7    349   VERMRDTVAGMRCVAGAIANIRCNLNGDMTPDFMAIYQDGQITLADEYQRTTMYNGFLGRLGFLPGRSREGERSMAERQI   428 (543)
Q Consensus    349   ~~~M~~t~~~~~~~~~~~~~~~~i~~~rc~lNG~~~PdFms~~~d~~i~l~D~~~r~~M~naflGrlG~~p~r~r~~~rs~aE~~i   428 (543)
                      ..+..+|......--.|.   -+++|=++++--.+-  ..+-+|=||-.+++--.-|..--|+...-.--.
T Consensus    338   ~~~~~~~t~~~sD~f~~I~---nisInfnn~~GLLSs--at~~~Ly~mSvkNGsnqswseFsG~~~~~~~~~~~---------   403 (508)
T 3J26_L      338   YQNLNNQITTPDVFLQIN---NLNLTWNNQQGILSG--ASSQNLYDFSVQNGYNKTWSEFNGVTQQFNGVS---------   403 (508)
T ss_dssp            HSSHHHHHSCCCCBCEEE---EEEEECTTCCCBTSS--SCHHHHHHHHHTTCCCSCTTTTTCEEEECCCST---------
T ss_pred             cccccccccCCCceeeee---eEEEEECCCcccCC--CCHHHHHHHHHHcCCCcchhheeeeeeccCCCC---------

Q ss_pred             HhcccCCCCcccccCCceeeEEEEEcCCCcceEEeecccccceeEEEEEechhhHhcC-CCCcEEEEEEEEhheeEEEe
Q Q_GrenLO7    429   LAYGGQGEGPERAEKPICSAGIVLTDPSMTLVREASSGALAGALSLQVQIAPQAQAAG-QVPSKLVVICITKQQIAFKK   506 (543)
Q Consensus    429   ~~y~~~~e~p~~~~~~l~~~~~~iv~TDPs~tvvReA~SG~l~GrlD~~i~f~~~~~~~~~~~~~~rL~vi~~~KqqIaf~k   506 (543)
                        +.+.     ....++-+||++-=..--=-.+..+-..-|...+||+.+-.-+... .....||||+..---+.|.+
T Consensus    404   -------------~~I~~~GSVl~L~pg~Dl~L~d~~a~gs~g~~~lQ~qv~~tN~~~~~~~~~~peL~Vi~~~~Gv~~~~~   474 (508)
T 3J26_L      404   ---GQPT-----KVIGLEGGIVCLELGKDVGLRDDEAEGVIGNFNLQVQMTVTNTNQYVTVTPDMYIVAVYDGTLVISN   474 (508)
T ss_dssp            ---TCCC-----EEEECCCCEECCTTTSSCCCSSCCSSCCCCCEEEEEEEEEECCCSSCCBCEEEEEEEEEEEEEEEEEEBT
T ss_pred             ---CCCc-----eEEeCCceEEEEeccCcccCCccccCcccceeEEEEEEEEEcCCCCCccCceEEEEEEecCEEEEeC
```

### Pink abalone xenomavirus

#### Replicase

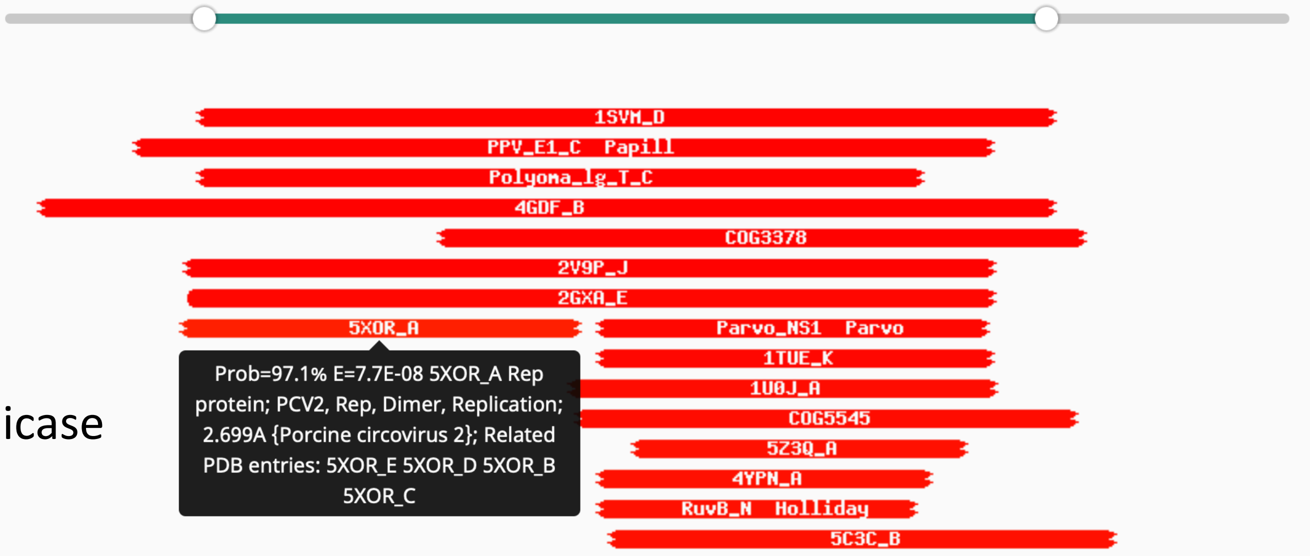


**5XOR_A Rep protein; PCV2, Rep, Dimer, Replication; 2.699A {Porcine circovirus 2}; Related PDB entries: 5XOR_E 5XOR_D 5XOR_B 5XOR_C**

Probability: 97.08%, E-value: 7.7e-8, Score: 91.84, Aligned cols: 129, Identities: 9%, Similarity: -0.131,

```
Q ss_pred              ecCCCCCCceeHHHHHHHHHHCCCCCC-cchhHHHhccCCCchhHccCCC---CchHHHHHHHccchhHHHcCcchHH-H
Q Q_3391286     81     VNQMRPDAKMNHQLLADWIIEREATSX-XYCNYRKFAREPVGEQMCQCGC---DYLVSIHKLHHDNAMAFANSPKVKE-V    155 (561)
Q Consensus     81     ~~~~~~~~~~~~~l~~n~~~~~~~t~~~i~~~y~~~~~~~~~~~~~~e~~~~~~~~h~h~~~~~~Na~~f~~ik~~~~~~    155 (561)
                       |....+.++++|++|++++     ++. ...+-......+.|+..+.|.+   |++|+|+|+.|+++++|.++++.++ .
T Consensus      6     ~~~~~~~~~~r~w~fT~~~~~~~~~~~~~~~l~~~~~~~~~~y~i~~~E~~~~tG~~HlQ~yv~f~~~~r~~~~k~~~~~    80 (158)
T 5XOR_A        6     NGRSGPQPHKRWVFTLNNP-----SEDERKKIRDLPISLFDYFIVGEEGNEEGRTPHLQGFANFVKKQTFNKVKWYLGAR    80 (158)
T ss_dssp              ------CCEEEEEEEEESC-----CHHHHHHHHTSBGGGEEEEEEEEEC---CCSCEEEEEEEEEEEEECHHHHHHHHCTT
T ss_pred              CCCCCCCCcCEEEEEecCC-----CHHHHHHHHhCCccccCEEEEEeeEeCCCCCcEEEEEEEECCCeeHHHHHHHHCCC


Q ss_pred              HHHHHHHHHHHHHHHhhcCceechhHHhccccCCCCCCCCeeeecCccccHHHHHHHHHHHHhHHHHhcCCCcchHHHHHH
Q Q_3391286    156     IKNGINQYRWIKQQVQENTTLAKTWSENFSNIGGDYTNKVLFPRNDFDRTYLFGRLKAIWGKFDSIFVKNDVIDVDRCYE    235 (561)
Q Consensus    156     ~~~~ie~~~g~~~qa~d~~y~~k~~~~~~~~~~~g~~~~~~~~~~~~~~~~~~~l~~~~~e~~~~~~e~f~~~~~~di~~~~~    235 (561)
                         .|+|.++|+..++.+  |+.|.                    ...          +.|+++.+..    |.++|+.++.+
T Consensus     81     --~h~e~~rgs~~~~~~~~Yc~Ke------------------~~~~~~~~~~~~~~~e~G~~~~q----G~r~dl~~~~~    124 (158)
T 5XOR_A       81     --CHIEKAKGTDQQNKE--YCSKE-------------------GNL---------LIECGAPRSQ----GQRSDLSTAVS    124 (158)
T ss_dssp              --CEEEECCSCHHHHHH--HHTTT-------------------SCE---------EEEEECCCC----------------
T ss_pred              --CccEecCCCHHHHHH--Hcccc-------------------CCe---------eEEcCCCCCC----CCCCcHHHHHH


Q ss_pred              HhhccCCchhhH-hchH
Q Q_3391286    236     DEYIDGFSKYVS-AGMA    251 (561)
Q Consensus    236     ~~~~~g~s~~~i-~~~~    251 (561)
                        .+..|.+..+| +..+
T Consensus    125     -~i~~g~~~~~I~~~~p    140 (158)
T 5XOR_A      125     -TLLESGSLVTVAEQHP    140 (158)
T ss_dssp              -----------------
T ss_pred              -HHHhCCCHHHHHHHCc
```

#### Penton

| # | Template | Alignment Coverage | 3D Model | Confidence | % i.d. | Template Information |
|---|---|---|---|---|---|---|
| 1 | c3iyjE_ | Alignment | 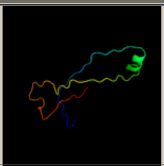 | 57.6 | 33 | **PDB header:**virus<br>**Chain:** E: **PDB Molecule:**major capsid protein l1;<br>**PDBTitle:** bovine papillomavirus type 1 outer capsid<br>Phyre2 Run Investigator |
| 2 | d1dzla_ | Alignment | 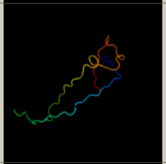 | 52.1 | 28 | **Fold:**Nucleoplasmin-like/VP (viral coat and capsid proteins)<br>**Superfamily:**Group I dsDNA viruses<br>**Family:**Papovaviridae-like VP<br>Phyre2 Run Investigator |
| 3 | c1dzlA_ | Alignment | 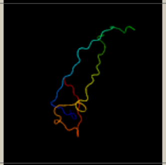 | 52.1 | 28 | **PDB header:**virus<br>**Chain:** A: **PDB Molecule:**late major capsid protein l1;<br>**PDBTitle:** l1 protein of human papillomavirus 16<br>Phyre2 Run Investigator |
| 4 | c6igdG_ | Alignment | 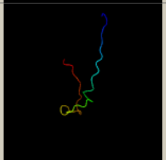 | 47.4 | 29 | **PDB header:**structural protein<br>**Chain:** G: **PDB Molecule:**major capsid protein l1;<br>**PDBTitle:** crystal structure of hpv58/33 chimeric l1 pentamer<br>Phyre2 Run Investigator |
