## Supplementary material for "Identification of Adomavirus Virion Proteins": Fig4Supp2 LO8 Tree

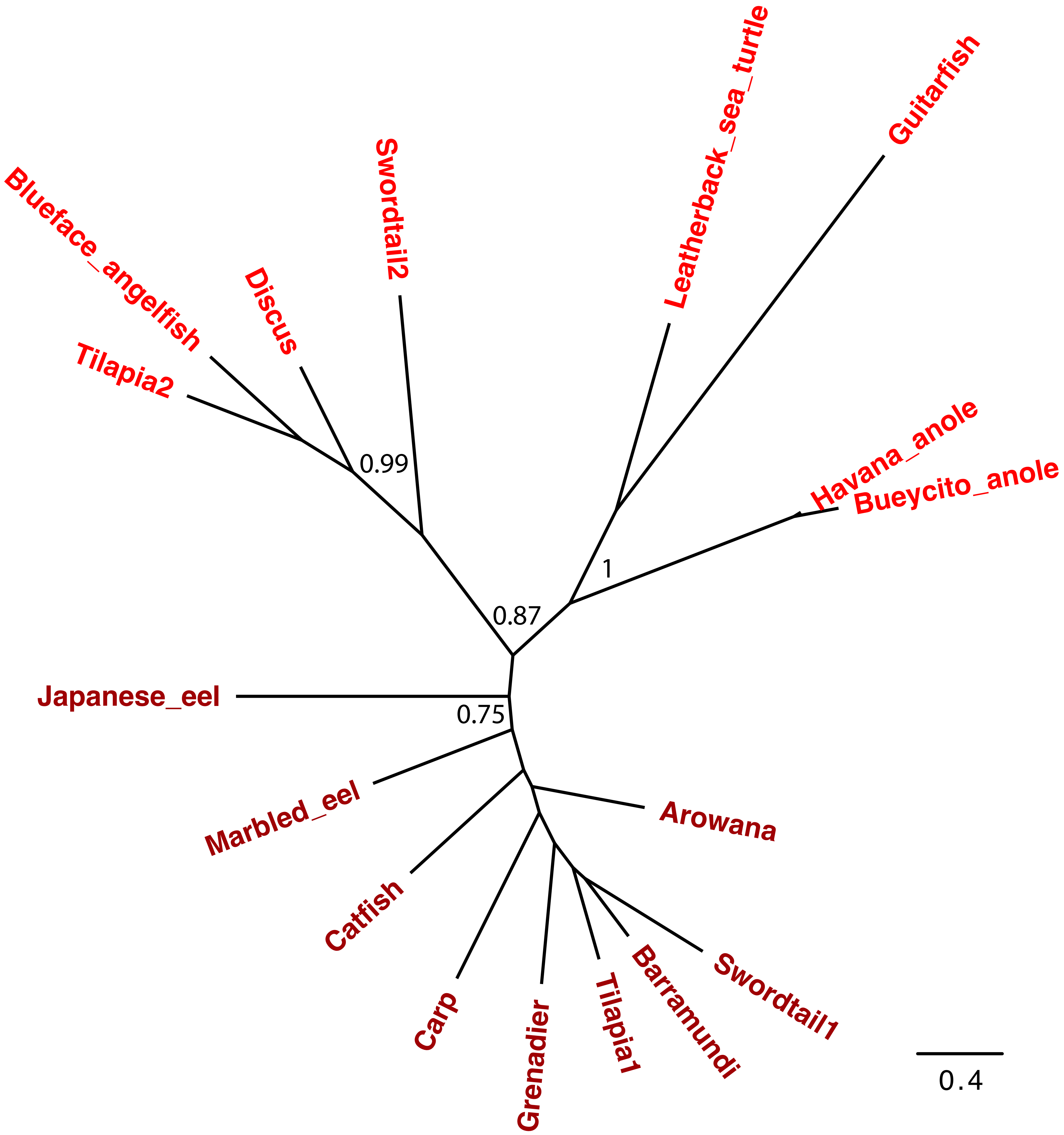


Figure 4 Figure supplement 2: A phylogenetic analysis of adomavirus LO8 (Adenain) proteins. Alpha adomavirus names are bright red, Beta adomavirus names are dark red. Bootstrap values are indicated for select nodes.
