## Supplementary material for "Identification of Adomavirus Virion Proteins": Fig7Supp1 VLP Electron Microscopy

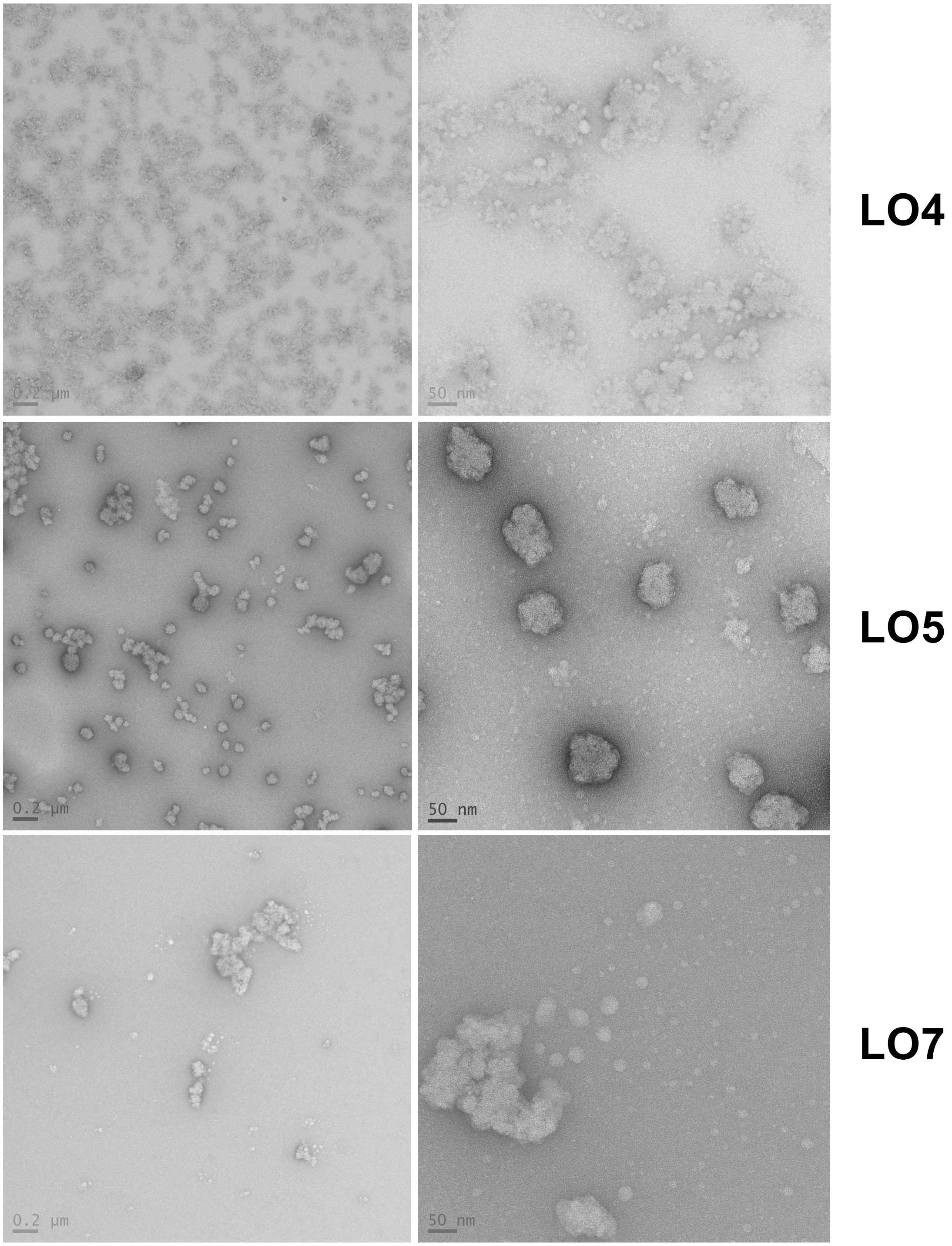


**Figure 7 Figure supplement 1: Negative-stain EM of particles formed after expression of single LO proteins in mammalian cells.** Mammalian 293TT cells were transfected with expression plasmids encoding the indicated marbled eel adomavirus proteins. Particles were extracted from cells by detergent lysis and purified by Optiprep ultracentrifugation. Scale bars are shown at the bottom left corner of each electron micrograph.
